# A latent cardiomyocyte regeneration potential in human heart disease

**DOI:** 10.1101/2023.09.14.557681

**Authors:** Wouter Derks, Julian Rode, Sofia Collin, Fabian Rost, Paula Heinke, Anjana Hariharan, Lauren Pickel, Irina Simonova, Enikő Lázár, Evan Graham, Ramadan Jashari, Michaela Andrä, Anders Jeppsson, Mehran Salehpour, Kanar Alkass, Henrik Druid, Christos P. Kyriakopoulos, Iosif Taleb, Thirupura S. Shankar, Craig H. Selzman, Hesham Sadek, Stefan Jovinge, Lutz Brusch, Jonas Frisén, Stavros Drakos, Olaf Bergmann

## Abstract

Cardiomyocytes in the adult human heart show a regenerative capacity, with an annual renewal rate around 0.5%. Whether this regenerative capacity of human cardiomyocytes is employed in heart failure has been controversial. Using retrospective ^14^C birth dating we analyzed cardiomyocyte renewal in patients with end-stage heart failure. We show that cardiomyocyte generation is minimal in end-stage heart failure patients at rates 18-50 times lower compared to the healthy heart. However, patients receiving left ventricle support device therapy, who showed significant functional and structural cardiac improvement, had a >6-fold increase in cardiomyocyte renewal relative to the healthy heart. Our findings reveal a substantial cardiomyocyte regeneration potential in human heart disease, which could be exploited therapeutically.

## Introduction

Loss of cardiomyocytes as a result of myocardial infarction or in the context of progressive cardiomyopathy can lead to heart failure, with considerable morbidity and mortality. Multiple strategies aiming to promote regeneration of the human myocardium are currently being explored. As cell replacement therapies using stem cell-derived cardiomyocytes still face considerable challenges (*1*), promotion of endogenous cardiomyocyte regeneration is an attractive alternative.

Cardiomyocytes in the healthy human heart have the capacity to renew for the lifetime of the individual, albeit at a low rate. In homeostasis, renewal occurs at a rate of around 0.5 % per year throughout adulthood, resulting in almost 40% of the ventricular cardiomyocytes being exchanged during life (*2, 3*). To what extent cardiomyocytes are regenerated in heart failure in humans is still poorly understood. Previous studies suggesting massive regeneration of cardiomyocytes proved difficult to reproduce (*4*), and identification of cardiomyocyte proliferation after a cardiac injury is challenging due to inflammation, scar formation, endoreplication and the proliferation of other cell populations (*5, 6*).

Depending on the underlying disease, cardiomyopathies are categorized as ischemic (ICM) or non-ischemic cardiomyopathy (NICM), with the latter comprising a heterogenous group of pathologies not primarily related to coronary artery disease. A left ventricular assist device (LVAD) is a pump that is surgically implanted in the left ventricle and helps propel blood, which is standard-of-care therapy for patients suffering from advanced heart failure and can be used as a bridge to heart transplantation or as a lifetime therapy. A subset of LVAD-supported patients attain significantly improved cardiac function and structure to the point where withdrawal of the LVAD support can be considered (*7–11*). However, the underlying mechanisms for LVAD-mediated myocardial recovery are not fully understood (*12*), and it remains unknown whether cardiomyocyte renewal contributes to this process (*6, 13, 14*).

In the present study, we retrospectively birth dated cardiomyocytes from patients with cardiomyopathy by measuring ^14^C, derived from nuclear bomb tests during the Cold War, in genomic DNA, a method we developed to study cell renewal dynamics in humans (*2, 3, 15*) (Supplementary Fig. S1). We established an integrated model of human cardiomyocyte renewal in cardiomyopathy with birth and death rates including endoreplication of cardiomyocytes. Our data demonstrate that cardiomyocyte renewal is minimal in failing hearts but can elevate well beyond the levels observed in healthy hearts through LVAD-mediated functional cardiac improvement.

## Results

### Cardiomyocyte nuclear ploidy and multinucleation increase in cardiomyopathy

Cardiomyocytes often exit the cell cycle prematurely and become polyploid without forming daughter cells. We determined nuclear ploidy in cardiomyocytes of patients with failing hearts diagnosed with NICM and ICM (**Supplementary Table S1)**. Nuclei were isolated from tissue samples and cardiomyocyte nuclei identified using flow cytometry by their positive staining for PCM-1 (21.3% ± 14.5% of all nuclei, Mean ± SD) (**Fig. 1A**), (**Supplementary Fig. S2B**) (*2, 16, 17*). Application of a DNA stain (DR) enabled the determination of their nuclear DNA content (**Fig. 1B**). We found a shift towards higher nuclear ploidy levels in both NICM and ICM **(Fig. 1C)** in comparison to healthy subjects (values taken from reference 2). As an integrated measure of nuclear ploidy, we define the nuclear ploidy level and set it to 100% for a pure diploid sample and 200% for a pure tetraploid sample. We observed a total increase in cardiomyocyte nuclear ploidy level from 176.9% [169.8%;196.9%] (median with interquartile range) in healthy hearts to 291.7% [250.4;341.4] in NICM and 221.1% [202.1%;255.0%] in ICM (**Fig. 1D**). The increase in nuclear ploidy was not linked to the age of the patient (NICM R=0.26, p=0.26; ICM R=0.49, p=0.11) (**Fig. 1E),** nor to the duration of disease (NICM R=0.19, p=0.55; ICM R=0.16, p=0.62) (**Supplementary Fig. S2D**).

**Figure 1.**
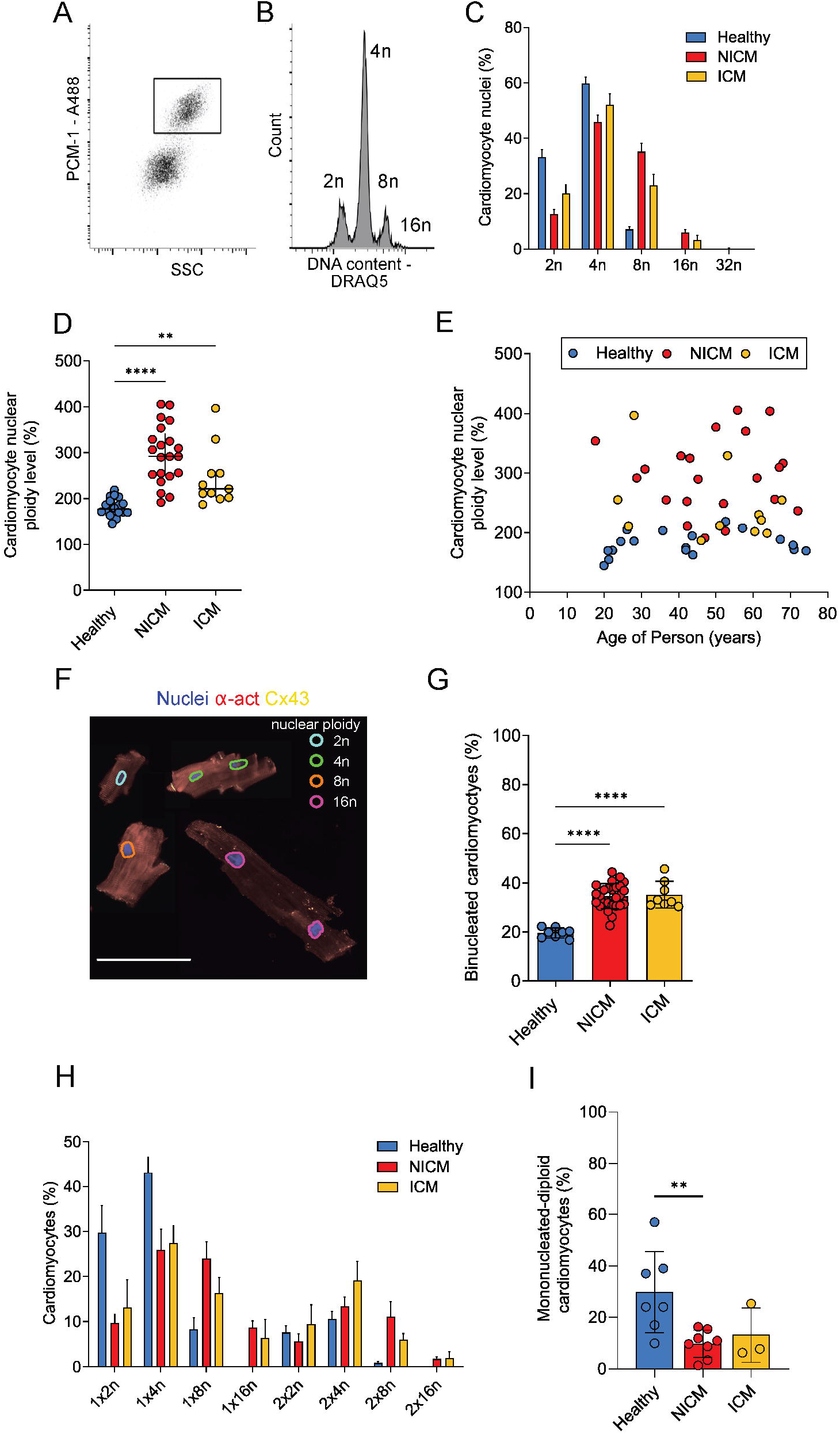
Isolation of human cardiomyocytes and ploidy assessment. (A) Flow cytometry-based sorting enables identification and isolation of cardiomyocyte nuclei with antibodies against PCM-1 (inlet). (B) Flow cytometry analysis of cardiomyocyte nuclei DNA content reveals their ploidy profile (2n = diploid, 4n = tetraploid, 8n = octaploid, 16n = hexadecaploid; shown is a representative NICM sample). (C) The distribution of cardiomyocyte nuclear ploidy populations shows a shift to higher ploidy levels in NICM (n=21) and ICM patients (n=11) compared to healthy adults (n=11, from (*2*)). (D) Cardiomyocyte nuclear ploidy level determined by flow cytometry is higher in both NICM (n=21) and ICM patients (n=11) than in healthy adults (n=11) (e.g. 100% corresponds to only diploid nuclei; 200% to only tetraploid nuclei) (Kruskal-Wallis one-way ANOVA on Ranks, H=30.3, p < 0.001, post hoc Dunn’s ** p=0.0063, **** p<0.0001), lines show median with interquartile range. (E) Ploidy increase in cardiomyopathy is not related to the age of the adult patient (NICM R=0.26, p=0.26; ICM R=0.49, p=0.11). (F) Image cytometry identifies cardiomyocyte ploidy classes. Cardiomyocytes were digested from tissue samples and stained for α-actinin (α-act) and connexin-43 (Cx43). The integrated intensity of the DNA dye was used to designate the nuclear ploidy class of each imaged nucleus, as shown by the overlay in different colors on the image. Cx43 enabled to determine if cardiomyocytes were intactly dissociated at the intercalated discs. Figure shows a compilation of several cardiomyocytes of different ploidy and nucleation levels. Scale bar size is 100 µm (G) The percentage of binucleated cardiomyocytes determined by image cytometry is higher in NICM (n=27) and ICM hearts (n=8) compared to healthy (n=8) (Ordinary one-way ANOVA, F=32.1; p < 0.0001, post hoc Tukey, **** p<0.0001). (H) Both binucleation and nuclear ploidy levels were determined in isolated cardiomyocytes from healthy (n=7) and pathological hearts (NICM n=8, ICM n=3) with image cytometry. (I) The percentage of mononucleated diploid cardiomyocytes is smaller in NICM 9.7% ± 5.2% and ICM 13.13% ± 10.56% compared to healthy hearts 29.7% ± 15.8% (Kruskal-Wallis one-way-ANOVA on ranks; H = 7.7; p = 0.021, ** p=0.0073).

In a second approach, we isolated intact single cardiomyocytes and performed image cytometry to determine the number of nuclei per cardiomyocyte and their ploidy status (**Fig. 1F, Supplementary Fig. S2G-H**). Using a sample set of healthy adults we found 19.6% ± 2.0% of cardiomyocytes to be binucleated (**Fig. 1G**), in agreement with previous studies, which also showed that this ratio remains unchanged in a healthy heart throughout the human lifespan (*2, 18, 19*). In contrast, we observed that cardiomyocytes isolated from NICM and ICM hearts showed significantly higher levels of binucleation compared to healthy hearts: 34.6% ± 5.1% in NICM and 35.2% ± 5.5% in ICM patients. Similarly, there was no correlation between the age of the patient and the ratio of binucleated cardiomyocytes (NICM R=0.13, p=0.52; ICM R=0.04, p=0.92;) (**Supplementary Fig. S2E**), nor with the disease duration (NICM R=0.01, p=0.96; ICM R=0.42, p=0.30) (**Supplementary Fig. S2F**).

Furthermore, we determined individual ploidy levels within mono- and binucleated cardiomyocytes. This analysis revealed that the proportion of mononucleated diploid cardiomyocytes (1x2n) is smaller in NICM (9.73% ± 5.2%) and ICM (13.13% ± 10.5%) compared to healthy hearts (29.7% ± 15.8%) (**Fig. 1H-I**). Although both nuclear ploidy and multinucleation increase in NICM and ICM, the distribution of ploidy levels within the mono- and binucleated cardiomyocytes remain similar in all sample groups (**Supplementary Fig. S2K**). These measurements by image cytometry are consistent with the extent of nuclear ploidy determined by flow cytometry, as shown by Bland–Altman comparison and Spearman correlation (**Supplementary Fig. S2I-J**). In conclusion, both methods demonstrate strongly increased cardiomyocyte DNA content in diseased failing human hearts from both ICM and NICM patients.

### 14C levels establish increased DNA synthesis in cardiomyocytes of failing hearts

We isolated cardiomyocyte nuclei (PCM-1 Positive) from NICM (n=16) and ICM patients (n=8), aged between 18 and 68, median age 51.5 (**Supplementary Fig. S2A)** by FACS (**Supplementary Fig. S2B-C**) and determined the genomic ^14^C concentrations by accelerator mass spectrometry (see **Methods, Supplementary Table S1, Fig. 2A**). The ^14^C amounts measured in NICM and ICM samples deviate more from the atmospheric ^14^C levels at the time of birth, than the values from healthy subjects (*2*), suggesting that there had been more cardiomyocyte DNA synthesis in these patients. To better classify this divergence, we determined the average ^14^C age of the cardiomyocytes’ genomic DNA (**Fig. 2B, Supplementary** Fig. 1A-B), which provides an initial estimation of cellular ages under the assumption that all cells were generated at the same time (*20*). In healthy subjects the genomic ^14^C age of cardiomyocyte DNA was on average 5.68 ± 1.22 years younger than the subject itself (**Fig. 2C**) (data set taken from reference 2). In contrast, we found the genomic ^14^C age of NICM cardiomyocytes to be 13.25 ± 5.44 years, and of ICM cardiomyocytes to be 11.32 ± 2.35 years younger than the individual, respectively. These observations demonstrate increased levels of DNA synthesis in cardiomyocytes from NICM and ICM patients compared to healthy controls.

**Figure 2.**
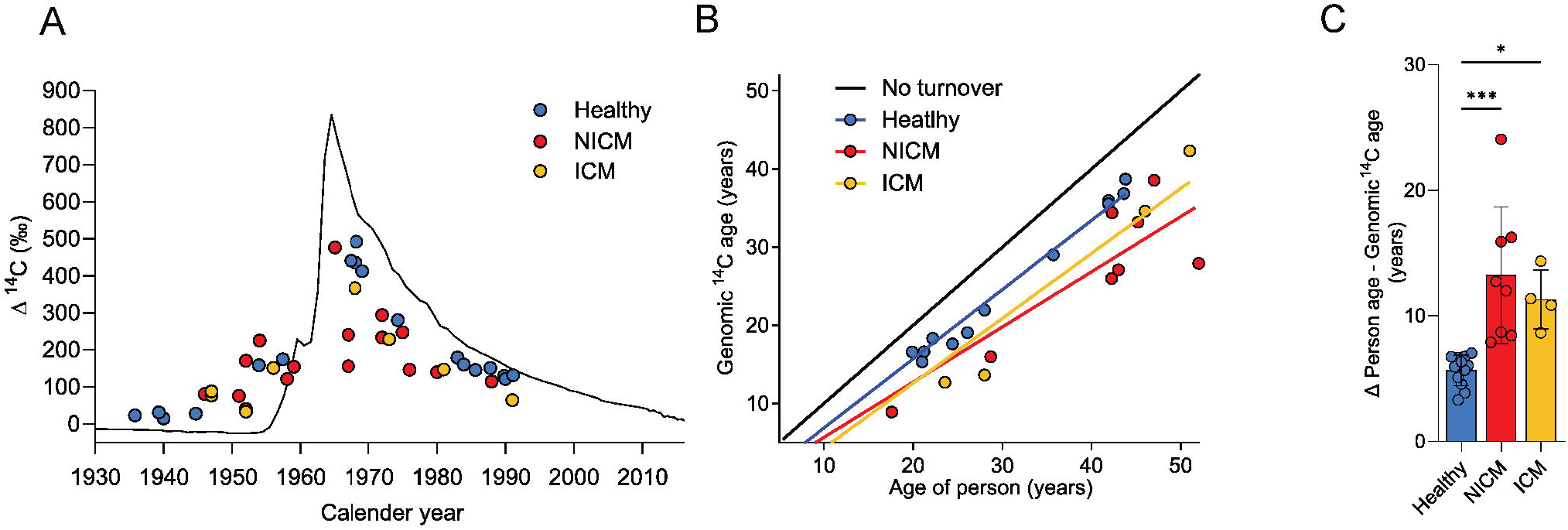
^14^C levels indicate increased DNA synthesis in cardiomyocytes of failing hearts. (A) Presentation of ^14^C data. The black curve indicates the historic atmospheric ^14^C concentrations. ^14^C measurements from heart samples are plotted as colored dots on the date of subject birth. Genomic ^14^C values of cardiomyocytes in healthy hearts (blue dots, n=18, data taken from Bergmann et al., 2015) are close to atmospheric values at the time of birth, indicating a limited postnatal and adult renewal of cells. The deviation of ^14^C values of cardiomyocytes in NICM (red dots, n=16) and ICM hearts (orange dots, n=8) from the atmospheric ^14^C curve suggest genomic DNA turnover. (B) The estimated genomic ^14^C age of cardiomyocytes was calculated from subjects with post-bomb birth dates and plotted at the person’s age. The black line indicates no turnover. (C) Both NICM (n=8) and ICM samples (n=4) show a higher deviation of the estimated genomic ^14^C age from the subject’s age compared to a healthy heart (n=12), suggesting increased DNA synthesis (Ordinary one-way ANOVA, F=12.38; p < 0.0002, post hoc Dunnet’s multiple comparison, *** p=0.0002, *p=0.017).

### Reduced cardiomyocyte renewal in heart failure

We defined a population balance equation for ^14^C concentration structured cell populations, which we used to predict ^14^C concentration dynamics from cell renewal rates (see mathematical methods) using a Bayesian inference framework (*20*). The cell renewal rates are allowed to change over time, which enables us to model different rates before disease onset, during the disease, and after treatment with LVAD. For NICM and ICM patients, we assumed that polyploidization and multinucleation proceed physiologically until disease onset and increased thereafter until tissue procurement. For this assumption, we applied the pathological ploidy levels individually measured for all ^14^C dated samples (**Fig. 1D**) and levels of binucleation determined for each population as depicted in **Fig. 1G**.

To establish the rate of cardiomyocyte renewal occurring specifically during the disease phase, we defined two phases of cardiomyocyte renewal: before and after the disease onset (first occurrence of heart failure symptoms). In this two-phase model, we assumed that the renewal rates before disease onset are the same as in healthy subjects (0.55% per year). The fitted renewal rates after disease onset resulted in much lower median renewal rate values of 0.03% per year for NICM patients and 0.01% per year for ICM patients, corresponding to a 18- and 50-fold reduction, respectively, compared to the healthy heart (**Fig. 3A**).

**Figure 3.**
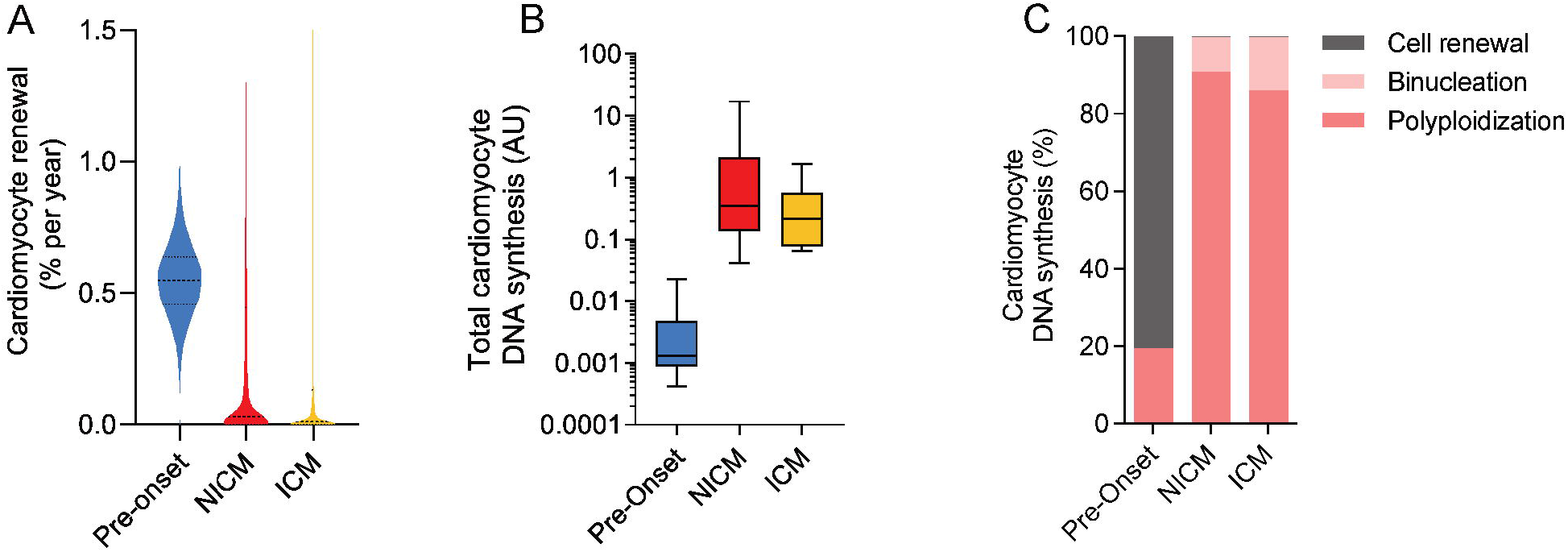
Mathematical modeling the renewal dynamics of cardiomyocytes in failing hearts. (A) With the two-phase renewal model the annual cardiomyocyte renewal rate can be determined before and after disease onset. In the disease phase, the percentage of newly formed cardiomyocytes drops from 0.55% [0.46%;0.64%] (median and interquartile range) to 0.03% [0.002%;0.45%] (median and interquartile range) in NICM, and 0.01% [0.001%;0.13%] (median and interquartile range) in ICM. (B) The annual amount of DNA synthesis modelled shows an increase in both NICM and ICM compared to healthy. (C) Percentage of DNA synthesis attributed to cell renewal, binucleation and nuclear polyploidization estimated in diseased hearts. After disease onset in NICM and ICM patients only < 0.3% of DNA synthesis can be attributed to proliferation-based cell renewal.

That cardiomyocyte generation drops greatly in heart failure, indicates that the increased ^14^C integration in cardiomyocyte DNA of failing hearts is almost exclusively due to polyploidization and multinucleation (**Fig. 2B-C**). To quantify this, we inferred total DNA synthesis from genomic ^14^C concentration in cardiomyocytes and examined the contribution of polyploidization, multinucleation and cardiomyocyte renewal due to proliferation to total DNA synthesis. Applying our two-phase mathematical model, we found that most of the DNA synthesis at the timepoint one year before disease onset was associated with proliferation-based cardiomyocyte renewal (80.6%) (**Fig. 3C**), whereas this rate dropped dramatically to values of 0.3% and 0.4% at timepoint of collection during the disease phase of NICM and ICM hearts, respectively. In those samples, most of the DNA synthesis was indeed associated with polyploidization (NICM 90.5%; ICM 87.2%) and multinucleation (NICM 9.2%; ICM 12.4%) (**Fig. 3C**).

In summary, polyploidization and multinucleation explain more than 99.5% of the ^14^C incorporated into failing cardiomyocytes’ DNA, while cardiomyocyte renewal drops drastically to minimal levels.

### Mechanical unloading and circulatory support increase cardiomyocyte renewal in patients with LVAD-mediated improvement of cardiac structure and function

Having observed the dramatically reduced cardiomyocyte renewal rate of the failing heart, we inquired whether this could be rescued by mechanical unloading and circulatory support. We analyzed advanced heart failure patients (including NICM and ICM) receiving LVAD therapy for 3 to 43 months (**Supplementary Fig. S4 A-B)**. Failing hearts not supported with an LVAD were used as the referent group (non-LVAD). It has been shown that in advanced heart failure patients, LVAD mechanical unloading and circulatory support can improve cardiac function to varying degrees (*8, 21*). Based on these LVAD studies as well as investigations of other heart failure therapies(*22, 23*), we categorized all patients with an absolute increase of left ventricular ejection fraction (LVEF) >5% as *responders*. To compare cardiomyocyte hypertrophy, we measured their cross-sectional area in these subgroups (**Supplementary Figure S4C**). Responders showed smaller cardiomyocyte cross-sectional area (832.8 ± 307 µm^2^) than patients with no improvement in LVEF (non-responders) (1123 ± 322 µm^2^) (**Supplementary Figure S4D**). Moreover, we studied mitochondrial abundance quantified by the relative copy number of human mitochondrial DNA by real-time PCR and normalized to genomic DNA copy numbers. Normalized mitochondrial DNA was lower in LVAD responders (1984 [1143;3736]) than in the LVAD non-responder group (4025 [2918;5233, p=0.02]), and non-LVAD patients (3487 [2821;5962, p=0.01]) (**Supplementary Fig. S4E**). Reversal of cardiomyocyte hypertrophy and decrease in mitochondrial load are two parameters of reverse remodeling that we have previously reported to be associated with mechanical unloading and circulatory support (*24*).

In contrast to previously published work (*13, 25*), we did not detect changes in nuclear ploidy or multinucleation in mechanically unloaded hearts using flow cytometry and image cytometry (**Supplementary Fig. S4F-G**). Neither responders nor non-responders showed significant differences in these parameters compared to non-LVAD advanced heart failure patients.

Next, we determined the ^14^C concentration in cardiomyocyte DNA in LVAD responders (n=15) and non-responders (n=13) (see Methods, Supplementary Table 1) to establish their ages and renewal rates in comparison to non-LVAD patients (n=24) (**Fig. 4A**). ^14^C concentrations of responders deviate more from atmospheric levels than non-LVAD patients. To classify potential fundamental differences in cardiomyocyte age within the three subgroups, we determined the genomic ^14^C ages as a first estimate and plotted these against the age of the patient (**Fig. 4B**). The group of responders was shown to differ the most from the no turnover prediction line. We compared these average genomic ^14^C DNA ages to the ages of the subjects and observed that it was 12.61 ± 4.61 years lower in the non-LVAD patients and 13.94 ± 3.65 years for non-responders, compared to 19.25 ± 6.04 years in responders (**Fig. 4C**).

**Figure 4.**
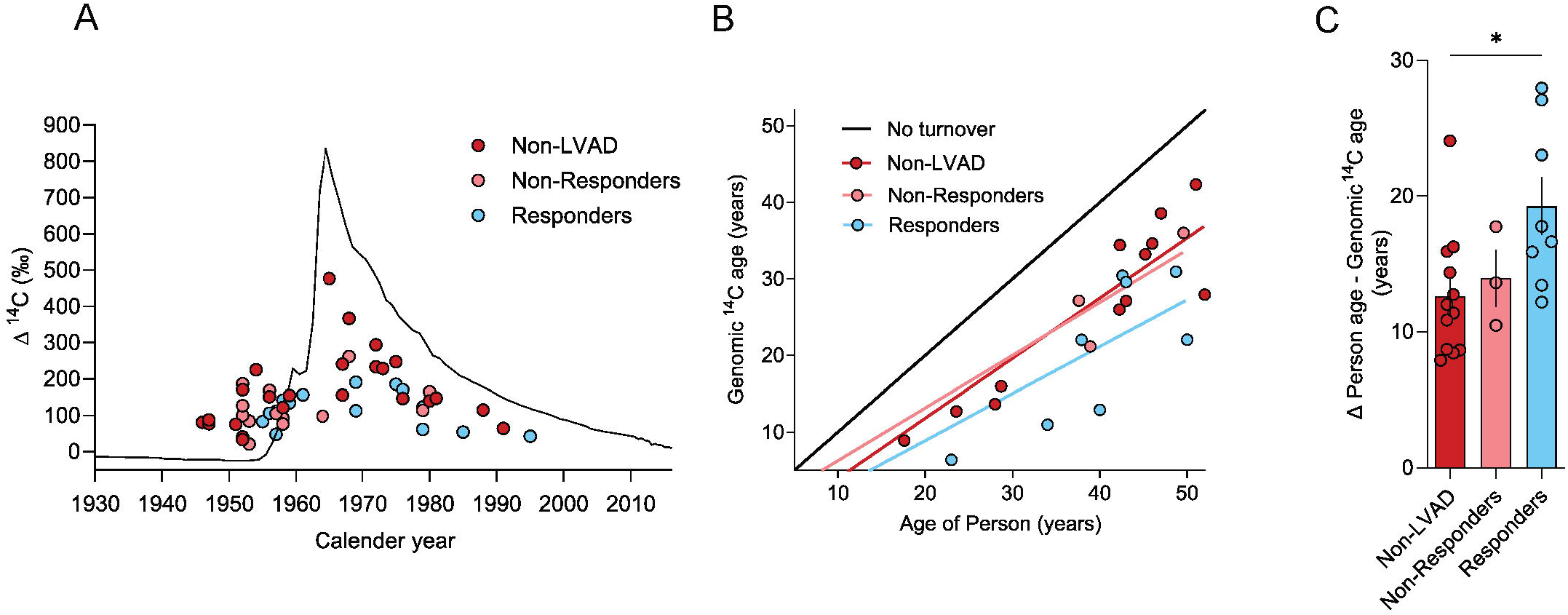
^14^C levels in cardiomyocytes of LVAD-supported hearts. (A) Presentation of ^14^C data from unloaded hearts. ^14^C measurements from heart samples are plotted as colored dots on the date of subject birth for diseased non-LVAD hearts (red dots, n=24, these correspond to NICM and ICM samples as shown in figure 2A), LVAD non-responders (n=13) and LVAD responders (n=15). The location of the dots from responder subjects suggests an increased deviation from the atmospheric ^14^C curve compared to non-LVAD subjects. (B) The estimated genomic ^14^C age of cardiomyocytes was calculated from subjects with post-bomb birth dates and plotted at the person’s age. The black line indicates no turnover. (C) Responders (n=8) show genomic 14C age which is on average 19.25 years younger than the person, compared to 13.94 years for LVAD non-responders (n=3) and 12.61 years for non-LVAD patients (n=12).

To further assess the impact of LVAD treatment on cardiomyocyte renewal during the disease process, we applied the above-described two-phase model of cardiomyocyte renewal. The fitted renewal rates after disease onset resulted in median renewal values of 0.03% per year for non-LVAD and 0.02% for LVAD non-responder patients; in contrast, renewal values in LVAD responders were 3.1% per year (**Fig. 5A**). In responders, 32.0% of all cardiomyocyte DNA synthesis can be attributed to cell renewal, in contrast to only 0.26% and 0.33% in non-responders and non-LVAD patients, respectively, in which almost all DNA synthesis is attributed to polyploidization and binucleation (**Fig. 5B**). This suggests a fundamental difference in the regenerative capacity in advanced heart failure patients exhibiting cardiac functional improvement during LVAD mechanical unloading.

**Figure 5.**
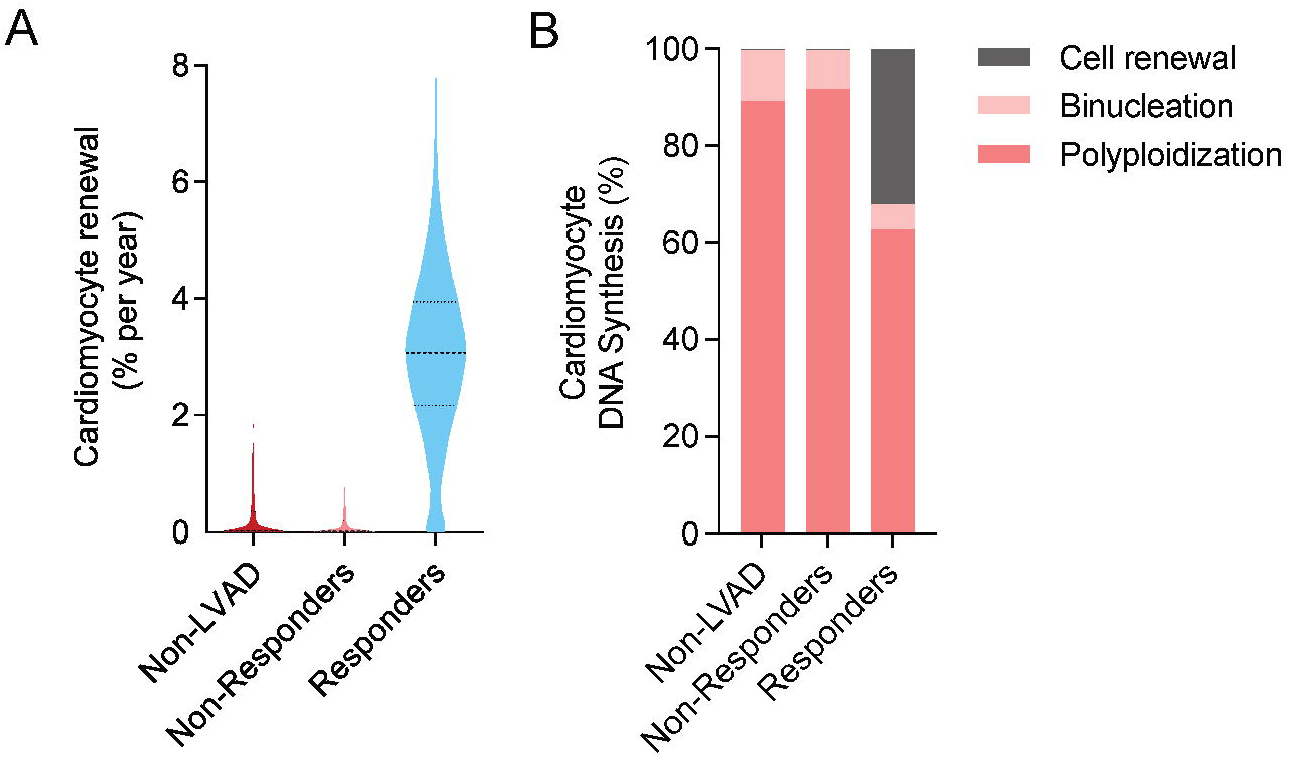
Mechanical unloading increases cardiomyocyte renewal in LVAD responders. (A) Cardiomyocyte renewal rate as determined using the two-phase renewal model. In cardiomyopathy, the percentage of newly formed cardiomyocytes per year is 3.07% [2.16%; 3.94] in LVAD responders compared to 0.03% in the non-LVAD group [0.002%;0.35%] and 0.02% in LVAD non-responders [0.001%;0.20%] (median and interquartile range). (B) Percentage of DNA synthesis attributed to cell renewal, binucleation and nuclear polyploidization determined at time of sample collection. In LVAD responders, 32.0% of all DNA synthesis can be attributed to proliferation-based cell renewal, in contrast to only 0.26% and 0.33% in non-responders and non-LVAD patients.

## Discussion

Here we have used retrospective ^14^C birth dating to investigate cardiomyocyte renewal in cardiac tissue samples from advanced heart failure patients. We report a major increase in DNA synthesis in all patient groups, leading mainly to cardiomyocyte polyploidization and binucleation. In patients without LVAD-mediated mechanical unloading, we found evidence of severely reduced cardiomyocyte generation compared to healthy subjects. In contrast, mechanically unloaded hearts exhibiting cardiac functional improvement (responders) showed a marked increase in cardiomyocyte renewal to levels approximately 6-fold higher than in healthy subjects. This demonstrates that there is a strong regenerative potential in the myocardium of a group of patients with advanced heart failure.

Our model predicts minimal cardiomyocyte renewal rates in both NICM and ICM patients (**Fig. 3A**), suggesting that cardiomyocyte renewal dynamics in failing hearts are not necessarily dependent on the underlying heart disease etiology. Even in patients with a ten-year history of cardiomyopathy, less than 0.2% of the existing cardiomyocytes would have been exchanged, which is much lower than in the healthy heart and unlikely to contribute to a clinical regenerative effect. The reduced capacity of cardiomyocytes to self-renew could even contribute to further disease progression.

Most studies on heart regeneration in both neonatal and adult rodents found an increase in cardiomyocyte proliferation directly after a myocardial lesion (*26, 27*). Newly formed cardiomyocytes generated mainly in the acute phase of the disease would have been detected with ^14^C birth dating unless these cells selectively died before ^14^C measurement. Thus, if these cardiomyocytes existed, they would be short-lived and have had a limited effect on regeneration. Given the chronic nature of heart failure, the minimal renewal rate is not surprising and could be a consequence of the hostile microenvironment present in failing hearts, which stimulates cell cycle activity but fails to accommodate successful cytokinesis (*28*).

Mechanical unloading and circulatory support have been shown to mediate structural, cellular, and molecular changes in the failing myocardium of advanced heart failure patients with different degrees of functional cardiac improvement (*8, 21*). Those patients showed an increase in cardiomyocyte renewal rate to 3.1% per year (**Fig. 5A**), which is approximately 6-fold higher than physiological cardiomyocyte renewal levels (*2*), along with reduced cardiomyocyte hypertrophy, and a lower mitochondrial load in these patients (**Supplementary** Fig. 4D-E) (*7, 24*). This has previously been associated with markers of cardiomyocyte proliferation, but that does not allow the assessment of whether new cells are generated and survive long-term, which we demonstrate here (*14*). Our results demonstrated that cardiomyocyte renewal is substantially elevated in the LVAD responder group, however, due to the limitation of our model, we cannot exclude the possibility that enhanced cardiomyocyte renewal is at least partly a pre-requisite for the reversal of heart failure observed in LVAD-supported hearts.

The discovery of a latent cardiomyocyte regenerative potential in the adult human heart identifies an attractive target for new therapies. This motivates studies to unveil the molecular regulation of this process, which would facilitate the development of pharmaceutical therapies for heart regeneration. For instance, mechanical unloading might reverse metabolic cascades that increase reactive oxygen species (ROS) production. This, in turn, can reduce oxidative DNA damage and activation of the DNA damage response (DDR) pathway (*29*) that causes cell cycle arrest in cardiomyocytes

(*30*). Indeed, we found a reduction of endoreplication and higher rates of cardiomyocyte proliferation in responders (**Fig. 5B**). This suggests that a successful approach for cell replacement strategies could be to selectively stimulate cytokinesis in already cycling cardiomyocytes (*31*).

In summary, our findings reveal a strong cardiomyocyte regenerative potential in advanced heart failure patients, which motivates studies focusing on the molecular mechanisms implicated in myocardial recovery and the discovery of pharmaceutical strategies to promote it.

## Acknowledgments

We thank M. Toro and S. Giatrellis for assistance and advice in flow cytometry, K. Håkansson for AMS sample preparation, G. Possnert for help with the analysis of the AMS samples, A. Felker and G. Eppens for help with flow cytometry sorting and DNA purification, and R. Savill for polyploidy analysis of isolated cardiomyocytes. This work was supported by the Flow Cytometry Facility, the Light Microscopy Facility, and the Histology Facility, all Core Facilities of the CMCB Technology Platform at TU Dresden. O.B. was supported by the Center for Regenerative Therapies Dresden, the Karolinska Institutet, the Swedish Research Council, the Ragnar Söderberg Foundation, the Åke Wiberg Foundation, and the LeDucq foundation. J.F. was supported by the Swedish Research council, the Swedish Cancer Foundation and Knut och Alice Wallenbergs Stiftelse. L.B. acknowledges support by the BMBF (grant 031L0293D). The model simulations and Bayesian inference were performed on HPC resources granted by the ZIH at TU Dresden. S.D. was supported by AHA 16SFRN29020000, NHLBI R01 HL 156667, NHLBI R01 HL135121, Merit Award I01 CX002291 U.S. Dpt of Veterans Affairs and the Nora Eccles Treadwell Foundation

The authors declare no competing interests.

---

**Supplementary Figure S1.** (A) Retrospective ^14^C birth dating can be used to estimate genomic DNA ages, utilizing the elevated atmospheric ^14^C concentrations caused by above-ground nuclear bomb testing in 1955–1963 (*15*). ^14^C in the atmosphere forms ^14^CO_2_ and is taken up by plants in photosynthesis. It reaches the human body through the food chain by consumption of plants or animals that live from plants. Thus, the ^14^C level in the human body parallels the atmospheric ^14^C concentration at all times. ^14^C is integrated into genomic DNA when a cell duplicates its chromosomes in S-phase, for example during cardiomyocyte renewal. As DNA is stable beyond cell division events, this creates a characteristic ^14^C datemark (*15*). To study cardiac cell renewal, nuclei are isolated from heart tissue samples and separated according to their cell type using FACS. Genomic DNA is extracted from the nuclei to determine its ^14^C concentration using accelerator mass spectrometry. The comparison of these measurements to historic atmospheric levels enables to establish cellular turnover dynamics by mathematical models.

(B) Method to determine genomic ^14^C age. Measured ^14^C data (red dot) is depicted at the person’s birthdate (1). The point of the atmospheric ^14^C curve (black line) with an equivalent ^14^C level (2, dashed line) would represent the time point of ^14^C formation of the analyzed cell population if all cells were born at the same time. This is the average birth date of the cardiomyocyte cell population. The difference between the time of sample acquisition and the time point of ^14^C formation (2) provides an estimate of the genomic ^14^C age and is used for the graphs in figure 2 B-C.

**Supplementary Figure S2**. (A) Age distribution of healthy subjects, patients with NICM and ICM as used for ^14^C birth dating. No difference in age distribution between the groups was found (ordinary one-way-ANOVA, p=0.368). (B) FACS gating strategy, (1) Nuclei were gated using SSC-A and FSC-A. (2) For doublet exclusion, nuclei were gated as singlets based on FSC-H/FSC-A. (3) Debris was excluded by DRAQ5-DNA staining and gating. (4) Next, PCM-1-FITC-A and SSC-A were used to discriminate PCM-1^+^ and PCM-1^-^ cells. (C) Sorted PCM-1^-^ (left panel) and PCM-1^+^ fractions (right panel) have high sorting purities as shown be re-analysis. (D) Duration of disease shows no correlation to nuclear ploidy level in NICM (R=0.19; p=0.55) and ICM patients (R= 0.16; p=0.62). (E) The degree of binucleation in cardiomyopathy is not related to the age of the adult patient (NICM R=0.13, p=0.52; ICM R=0.04, p=0.92). (F) Duration of disease shows no correlation to fraction of binucleation in NICM and ICM patients (NICM R=0.01, p=0.96; ICM R=0.42, p=0.30). (G-H) Example histograms of integrated intensity (the sum of the pixel intensity over all of the pixels in an object) of measured nuclei in isolated cardiomyocytes from healthy subject and NICM patient. Based on the peaks in the histogram ploidy classes can be attributed to each nucleus. (I) Bland-Altman comparison to FACS analysis verifies the accuracy of the image cytometry-based determination of nuclear ploidy. (J) Nuclear ploidy determined via image cytometry and flow cytometry show a strong correlation, R= 0.77, p=0.0012. (K) The nuclear ploidy distribution is largely similar between mono- and binucleated cardiomyocytes. Bars are shown as mean with SEM.

**Supplementary Figure S3.**

(A and B) Schematic visualization of two different model scenarios (I and II) describing the birth and death of cardiomyocyte cell populations with different ploidy levels. (C) To validate these scenarios, we fitted previously generated ^14^C data from healthy hearts reported by Bergmann et al. 2015. Both scenarios obtained similar renewal rates of 0.58% [0.49%;0.67%] (median and interquartile range) (Scenario I) and 0.55% [0.46%;0.64%] (median and interquartile range) (Scenario II) per year.

**Supplementary Figure S4.**

(A) Age distribution of non-LVAD and LVAD patients as used for ^14^C birth dating. (B) Distribution of LVAD therapy duration in non-responders and responders as used for ^14^C birth dating. (C) To assess cardiomyocyte cross-sectional area (CSA) cell borders were visualized in myocardial sections, which were sectioned perpendicular to the fibers, using a wheat germ agglutin (WGA) staining. CSA was determined using a CellProfiler pipeline which excluded objects such as vessels based on size and shape. (D) Cardiomyocyte CSA was found to be reduced significantly in responders (n=22) compared to patients with no improvement in EF (non-responders, n=11) (One-way-ANOVA P=0.03; F=3.75; post hoc Holm-Sidak against responders, *p=0.03), bars show median with interquartile range. (E) Normalized mtDNA was significantly lower in LVAD responders (n=20) than in the LVAD non-responder group (n=15), and non-LVAD patients (n=21) (Kruskal-Wallis One-way-ANOVA on ranks, P=0.009; H=9.4; post hoc Dunn’s, *p=0.02, **p=0.01). (F) Cardiomyocyte nuclear ploidy level does not differ between non-LVAD (n=41), non-responders (n=14) and responders (n=20) (Kruskal-Wallis One-way-ANOVA on ranks H=4.1; P=0.13). (G) The percentage of binucleated cardiomyocytes is similar in non-LVAD patients (n=26), LVAD non-responders (n=8) and responders (n=14) (Ordinary One-way ANOVA, F=1.64; p = 0.20).

**Supplementary Table 1.**
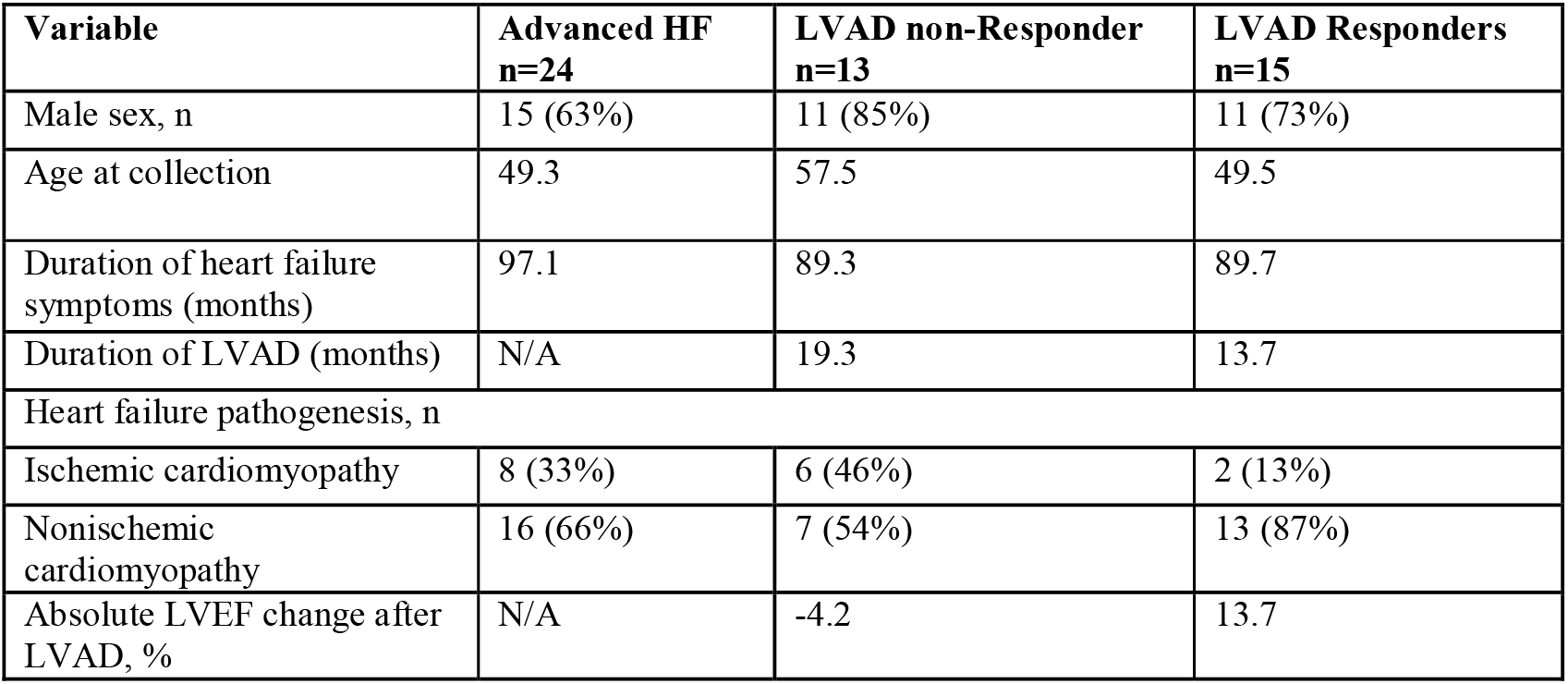
Characteristics of patients included in ^14^C study.

**Supplementary Table 2.** List of analyzed human subjects and heart tissue. Description of study subjects including diagnoses of heart disease and measured ^14^C values, and ploidy levels. NICM = non-ischemic cardiomyopathy; ICM = ischemic cardiomyopathy; ND = not determined

