## Supplementary figures and images for "A latent cardiomyocyte regeneration potential in human heart disease"

### Supplemental Figures

A

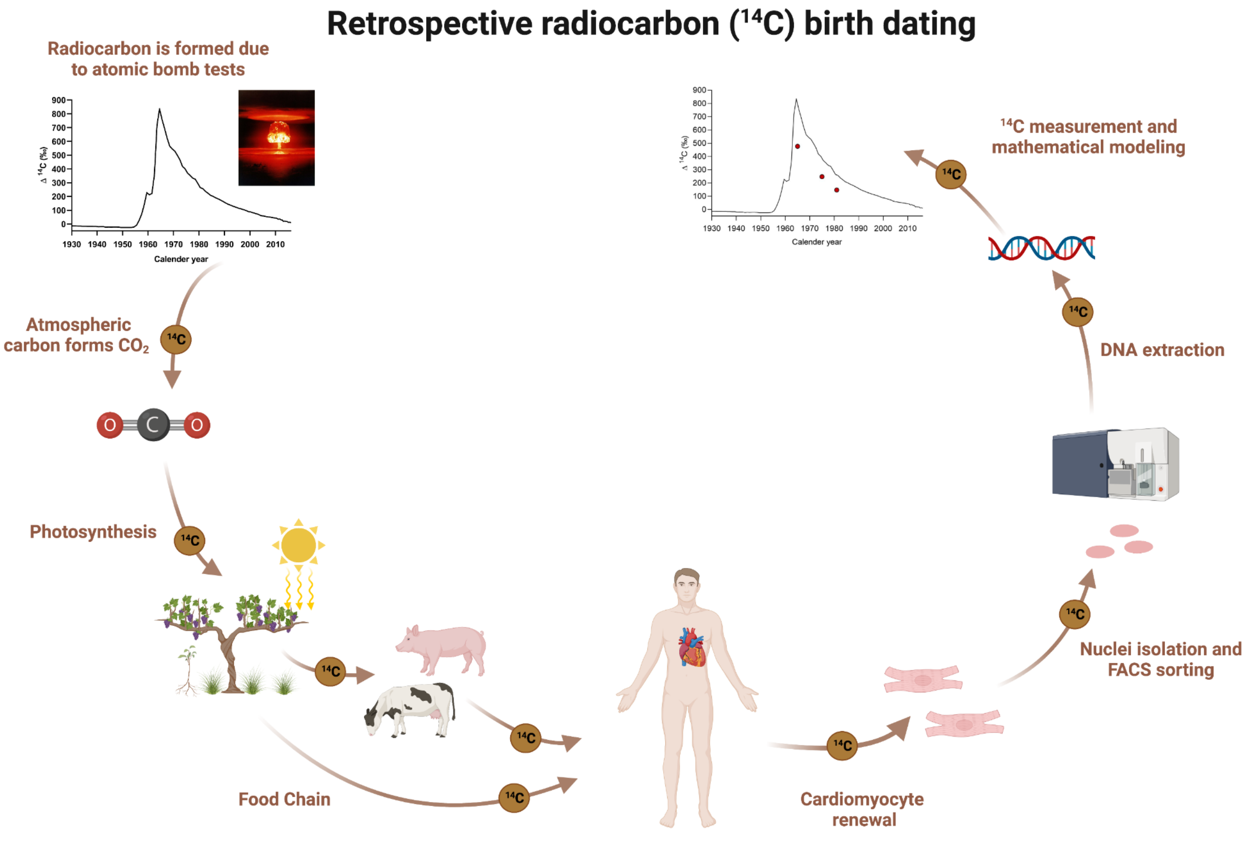


B

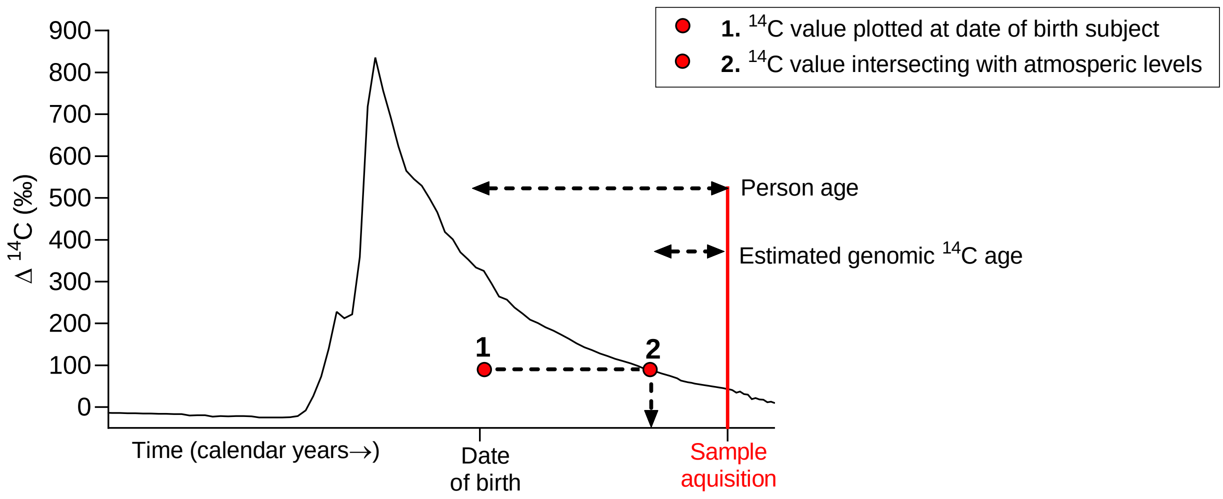


Supplemental Figure 1

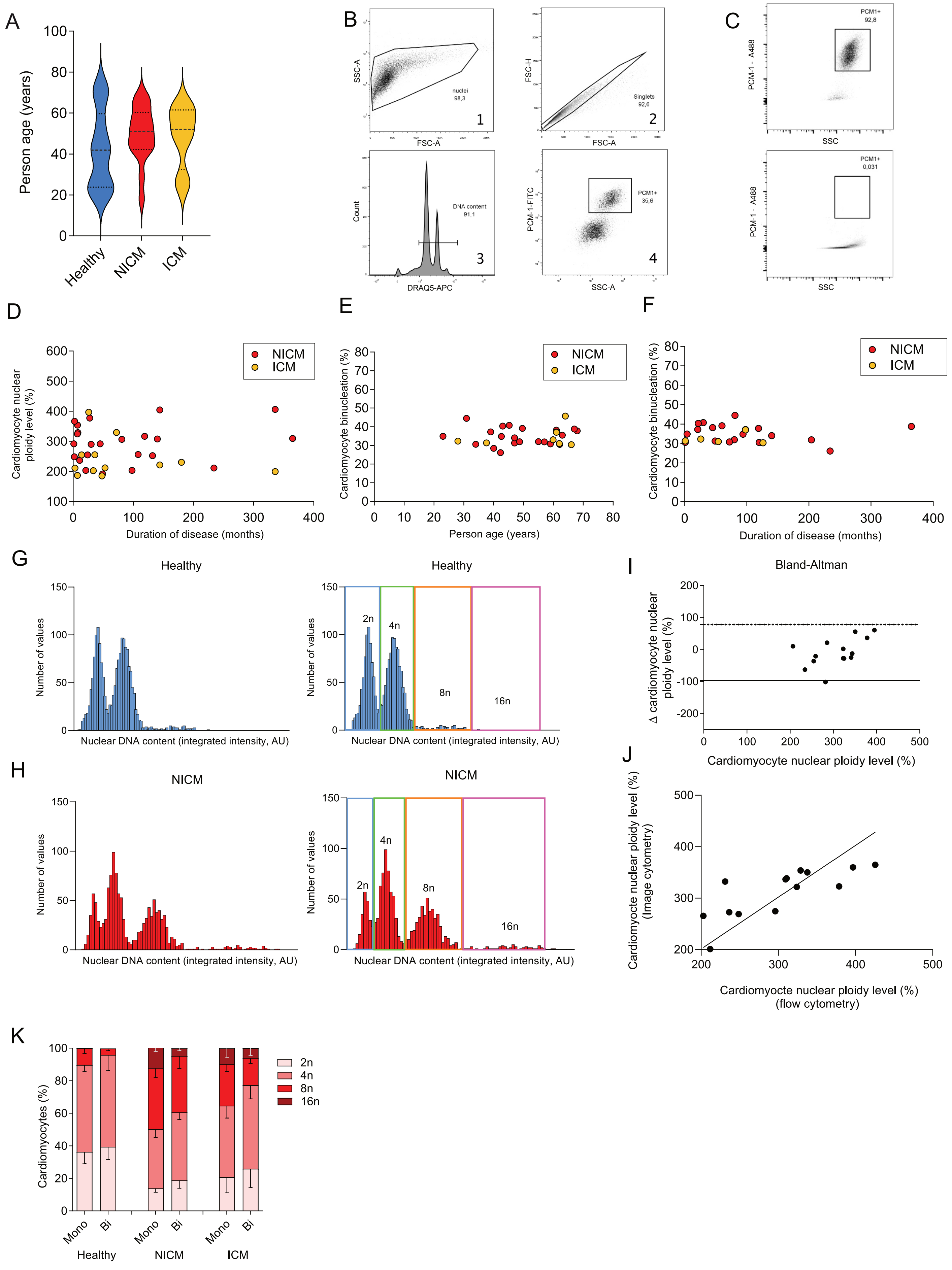


Supplemental Figure 2

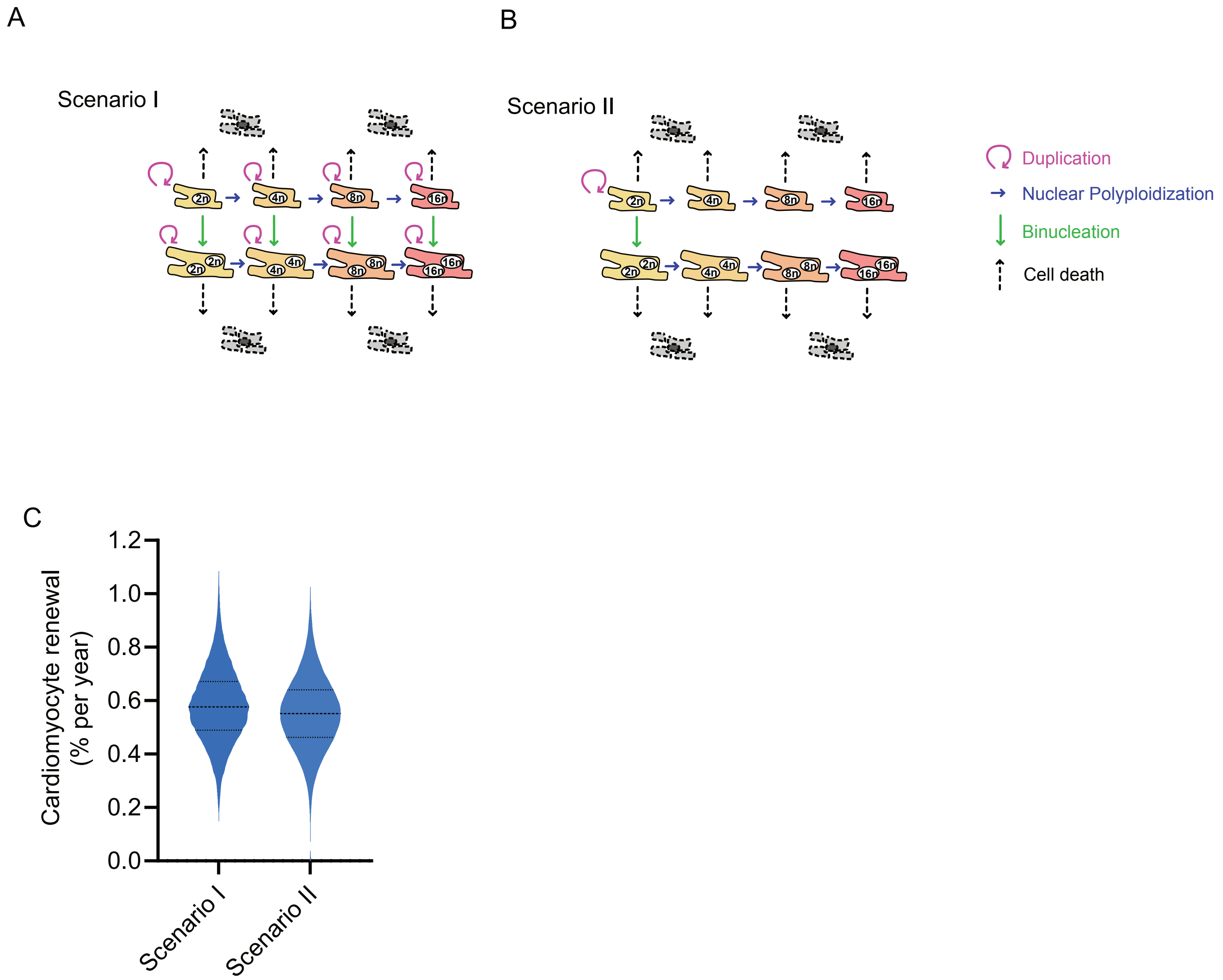


Supplemental Figure 3

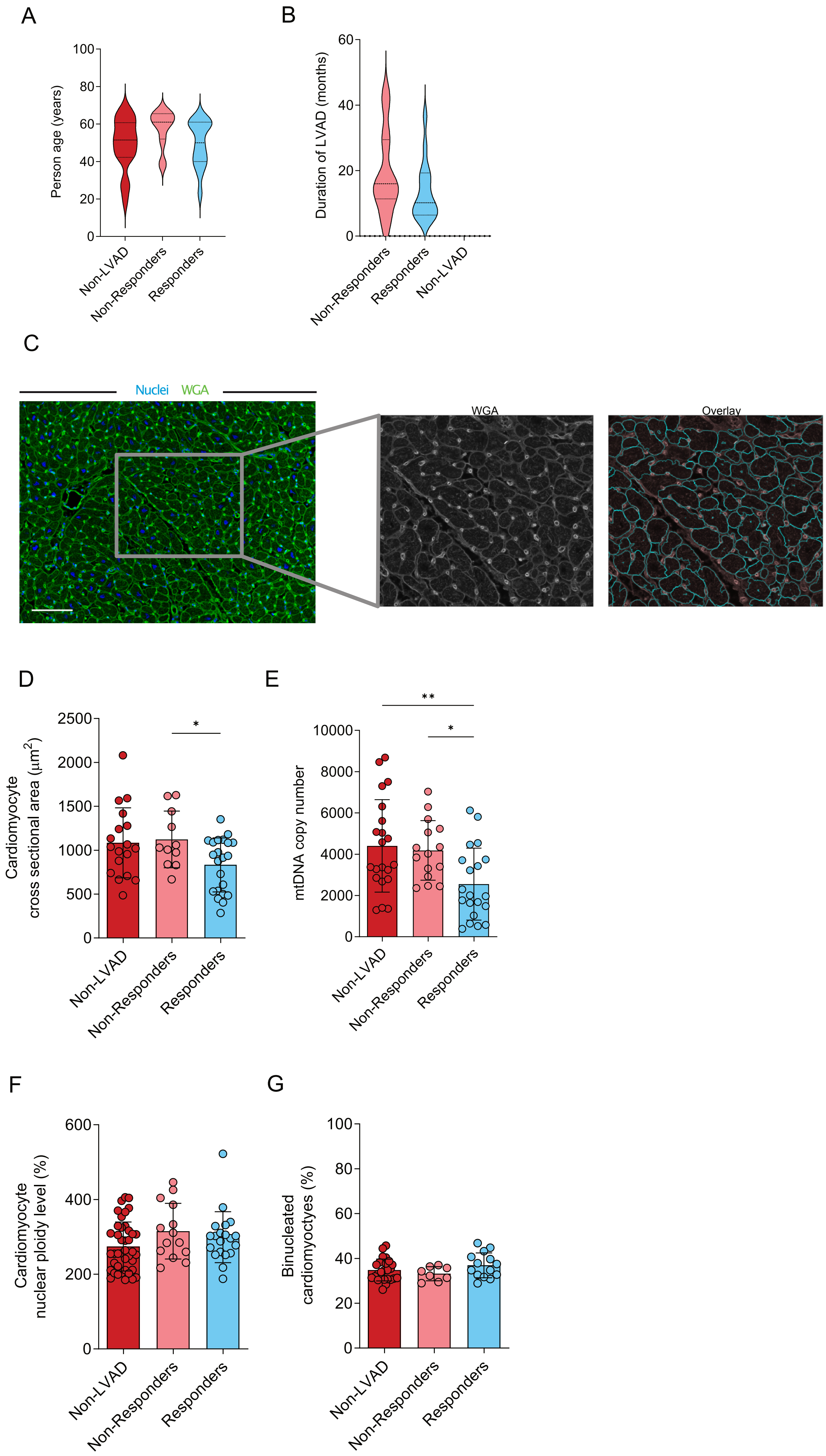


Supplemental Figure 4
