## Supplemental Methods for "A latent cardiomyocyte regeneration potential in human heart disease"

### Supplementary Methods

#### Tissue collection and preparation

Heart tissue was procured from The European Homograft Bank in Brussels, Belgium; Medical University of Graz, Austria; KI Donatum, Karolinska Institute Stockholm, Sweden; University of Lund, Sweden; Institutions comprising the Utah Transplantation Affiliated Hospitals (U.T.A.H.) Cardiac Transplant Program (ie, University of Utah Health Science Center, Intermountain Medical Center and the Veterans Administration Salt Lake City Health Care System), Spectrum Health Universal Biorepository, Utah, USA, and Gift of Life, Michigan, USA. Ethical permissions have been obtained from local ethical committees. Relevant clinical parameters are listed in Supplementary Table 1. Tissue from the left ventricle and septum harvested at the time of heart transplantation, and tissue from the left ventricle apical cores removed during LVAD implantation were collected, dissected, and stored at -80°C until further processing.

#### Cardiomyocyte nuclei isolation

Frozen heart tissue (approx. 10 g) was dissected and dissociated in a blender in lysis buffer (0.32 M sucrose, 10 mM Tris-HCl pH = 8.0, 5 mM $CaCl_2$, 5 mM MgAc, 2 mM EDTA) for 10 min. The tissue was further homogenized with a T-25 Ultra-Turrax probe homogenizer (IKA Germany) at 24,000 rpm for 10 sec and afterward dounced with a type A pestle in a 40 mL glass douncer (VWR), applying 8 strokes. The nuclear isolate was filtered through 100 µm and 60 µm nylon mesh cell strainers and sedimented at 500 × g for 10 min. The pellet was dissolved in sucrose buffer (2.0 M sucrose, 10 mM Tris-HCl pH 8, 5 mM MgAc), and layered onto 10 ml sucrose buffer in a BSA-coated centrifuge tube. The samples were spun down at 26,500 × g for 60 min in a Beckman Avanti Centrifuge (Beckman Coulter). The nuclei pellets were dissolved in NSB buffer (0.44 M sucrose, 10 mM Tris-HCl pH 8.0, 70 mM KCl, 10 mM $MgCl_2$, 2 mM EDTA). All steps were performed at

4°C.

### Immunolabeling and flow cytometry

A polyclonal rabbit anti-Pericentriolar material 1 (PCM-1, Sigma Aldrich, HPA023374) antibody, conjugated to Alexa Fluor® 488 or without conjugation to a fluorochrome, was added to the sample in a 1:800 dilution and incubated overnight. The following day the samples were filtered through 40 µm cell strainers (BD Bioscience) and the sample was labeled with the DNA stain DRAQ5® (ThermoFisher, 62251, 1:1000) for ploidy analysis, and when processed without conjugated primary antibody with secondary anti-rabbit Alexa Flour® 488 (1:1000) for 1 hour. Subsequently, the sample was loaded to the flow cytometer (BD FACS Aria II), single nuclei were identified by FSC-H/FSC-A gating, and cardiomyocyte nuclei were defined by gating the PCM-1 positive nuclei. After nuclei sorting, the populations were reanalyzed for purity. The sorted nuclei were spun down at 500 × g for 10 min, dissolved in PBS, and kept at -20°C until further processing. All steps were performed at 4°C. If the sorting purity was less than 100%, the DNA purity of the sample was calculated according to the following equation:

$$Purity_x = \frac{f_x * c_x}{f_1 * c_1 + f_2 * c_2 + \ldots + f_n * c_n}$$

Where $x$ is the sample being corrected, $f_x$ is the nuclei purity measured in the FACS, $f_{1 \ldots n}$ are the fractions of the $n$ contaminating populations, $c_{1 \ldots n}$ are the DNA contents of the contaminating nuclei (e.g. 4n: $c_1$=4), reflecting that polyploidy nuclei contain more DNA.

### DNA extraction for $^{14}C$ birth dating

DNA extraction was performed under clean room conditions. 500 µl DNA lysis buffer (1%

SDS, 5 mM EDTA-$Na_2$, 10 mM Tris-HCl pH 8) and 6 µl Proteinase K (20 mg/mL Life Technologies, EO0491) were added to each sample. The samples were incubated at 64°C overnight. After adding -half the volume of 5 M NaCl, the samples were vortexed for 15 sec and spun down at 16,000 × g for 3 min. The supernatants were transferred to 10 ml glass tubes, 3x volumes of 96% ethanol were added, and the glass tube was inverted several times until a DNA precipitate became visible. The DNA pellet was washed in DNA washing buffer (70% ethanol, 0.5 M NaCl) 3 times for 15 min each. The washed DNA pellets were dissolved in 200 µl sterile double distilled water (Gibco) and allowed to dry at 64°C overnight. The dried DNA pellet was dissolved in 500 µl sterile water overnight. The sample was analyzed for DNA concentration and purity by measuring the absorbance at 260 nm and 280 nm with a spectrophotometer (NanoDrop 1000, Thermo Scientific).

**Accelerator Mass Spectrometry**

Accelerator Mass Spectrometry measurements were performed blind to the identity of the samples and as described previously (*32*). Purified DNA samples were lyophilized to dryness. Excess CuO was added to each dry sample and the tubes were evacuated to $2.10^{-3}$ mbar and sealed with a high-temperature torch. The tubes were placed in a furnace set at 900°C for 3.5 h to combust all carbon to $CO_2$. The $CO_2$ was then cryogenically purified, trapped, and reduced to graphite in the presence of an iron catalyst in individual reactors at 550°C for 6 hours. Graphite targets were measured at the Department of Physics and Astronomy, Ion Physics, Uppsala University (*33*). Stable Isotope ratio, $\delta^{13}C$, was measured for each sample to correct for any isotopic fractionation. Strict laboratory practice was implemented to minimize stray background carbon contaminating the samples. Corrections for background carbon contamination introduced during sample preparation were made as described (Hua et al., 2004; Santos et al., 2007). $^{14}C$ data is reported as decay-corrected $\Delta^{14}C$. The measurement accuracy was in the range of 1 to 3 % (2 $\sigma$).

**Cardiac sectioning and immunostaining**

Wheat Germ Agglutinin (WGA) staining was performed on 5 μm paraffin sections prepared from the formaldehyde fixed tissue samples. Sections were deparaffinized using xylene and ethanol and incubated with Wheat Germ Agglutinin, Alexa Fluor™ 488 Conjugate (1:1000, ThermoFisher, W11261) diluted in PBS. Sections were washed three times with PBS and mounted with ProLong Gold Antifade Reagent containing DAPI (Life Technologies, P36935). Cardiomyocyte cross-sectional area was determined using CellProfiler 2.0 (*34*), analysis pipeline is provided as a data file (https://zenodo.org/record/7560219).

**mtDNA counts**

To analyze mtDNA content total DNA was extracted from the tissue samples for processing with Takara Human mtDNA Monitoring Primer Set (Cat #7246). Tissue biopsies were taken before the start of the nuclei isolation procedure, using a 6 mm wide disposable biopsy punch (Kai medical, #48601) and homogenized by the Ultra-Turrax® homogenizer at 24,000 rpm for 10 seconds in 1 mL lysis buffer (0.32 M sucrose, 5 mM CaCl2, 5 mM magnesium acetate, 2 mM EDTA and 10 mMTris-HCl, pH 8.0). DNA was isolated using 200 µL of the lysate, with the QIAamp® DNA Blood Mini Kit (Qiagen, 51104) according to the manufacturer's instructions. DNA concentration and purity was determined using NanoDrop. Quantitative real-time PCR reactions were set up in a total volume of 25 µL using the Takara Human mtDNA Monitoring Primer Set (Takara, 7246), to quantify the relative number of copies of human mtDNA with respect to the nuclear genomic DNA (nDNA). The set contains primer pairs for the amplification of four regions: two primer pairs for the detection of mtDNA (ND1 and ND5) and two primer pairs for the detection of nDNA (SLCO2B1 and SERPINA1). Reactions were setup following manufacturers protocol using 12.5

µL of SYBR® Green master mix (Thermo Fisher, #4309155) and 10 ng of template DNA. Thermocycling was set up in the StepOne™ Real-Time PCR System (Applied Biosystems, #4376357).

**Cardiomyocyte isolation**

Tissue samples were dissected into 0.5 – 1 mm cubes and fixed in 3.7% formaldehyde solution for 100 min at RT. Subsequently, tissue blocks were washed three times with PBS for 30 min and then digested in a digestion buffer consisting of collagenase B (Roche, 11088815001) (3.6 mg/mL) and collagenase D (Roche, 11088858001) (4.8 mg/mL) in PBS for 24h at 37°C with gentle rotation. Isolated cardiomyocytes were gently centrifuged at 1 x g and the supernatant was removed before resuspending in PBS. For several samples, this procedure was repeated for an additional 24h on the remaining tissue.

**Immunostaining of digested myocytes**

Isolated cardiomyocytes were allowed to settle for 15 min and were resuspended in 500 µL blocking buffer with primary antibodies Mouse-α-Actinin,1:500 (Sigma Aldrich, A7811) and Rabbit-Connexin-43, 1:1000 (Sigma Aldrich, C6219) for 30 min. Cardiomyocytes were washed once with PBS for 15 min and incubated with secondary antibody solution Alexa Fluor™ 555 anti-mouse, 1:1000 (Jackson Immuno Research, 711-546-152) and Alexa Fluor™ 488 anti-rabbit, 1:1000 (Abcam, ab150110) and Hoechst 33342, 1:1000 (ThermoFisher, 62249) for 30 min. Cells were washed with PBS and resuspended in a small volume of PBS and pipetted onto a glass slide. Cardiomyocytes on the glass slide were allowed to settle and adhere before mounting (ProLongTM Gold antifade reagent, ThermoFisher, P36930). All imaging was performed on a Keyence BZ-X800E using 20x objectives with appropriate dichroic filters. Exposure times were chosen that allowed capturing of the full dynamic range without overexposure and were similar for all acquired images. At least 30 fields of views were recorded per sample in a non-overlapping manner.

**Quantification of ploidy and binucleation on sections**

To identify nuclei and measure their integrated intensities a CellProfiler 4.0 (*35*) pipeline was set up (https://zenodo.org/record/7560219). Nuclei identification was performed using the "identify primary objects" module with an adaptive thresholding method to account for background variances. The α-Actinin image was used to exclude non-cardiomyocyte nuclei that do not fall within the α-Actinin positive areas. Further filtering by several neighbors and solidity was chosen to exclude large clumps of nuclear stains and irregular shapes, which indicated bad segmentation or accumulation of staining that did not resemble individual nuclei. Area and shape descriptors as well as intensity descriptors were exported for further analysis and integrated fluorescence intensity of the Hoechst stain was plotted as histograms to determine ploidy thresholds manually. These thresholds were then used to overlay colored outlines on the overlay pictures for further analysis. Evaluation of the ploidy levels in mono- and binucleated cells was performed manually based on the overlay pictures, considering cardiomyocyte shape and border morphology using connexin-43 staining.

**Clinical Procedures**

**Echocardiography**

Twenty eight out of 52 advanced HF patients, requiring circulatory support with continuous flow LVAD were prospectively enrolled at 1 of the institutions comprising the Utah Transplantation Affiliated Hospitals (U.T.A.H.) Cardiac Transplant Program (ie, University of Utah Health Science Center, Intermountain Medical Center and the Veterans Administration Salt Lake City Health Care

System). Patients who required LVAD support because of acute HF (acute myocardial infarction, acute myocarditis, postcardiotomy cardiogenic shock, etc) were excluded. Serial echocardiographic assessments were performed prospectively within 1 month prior to LVAD implantation and at 1, 2, 3, 6, 9, 12 post LVAD implantation based on a protocol that has been previously described (PMID: 23500219). Patients were categorized as responders and non-responders based on the absolute LVEF change pre to post LVAD implantation. Patients with an absolute LVEF increase > 5% within 1 year on LVAD support were defined as responders.

**Statistics**

Values are given as Mean ± SD unless stated otherwise. Statistical tests were performed, and graphs were generated using Graphpad Prism Version 9.5.0.

**Mathematical modeling of cardiomyocyte renewal**

In the following, we outline the mathematical models, describe the methods for parameter estimation and model selection, and present the results based on the models. Our aim was to devise a mathematical model that predicts the genomic $^{14}C$ concentration dynamics based on cellular turnover rates. This enabled us to estimate cellular turnover rates by fitting the model predictions to the measured genomic $^{14}C$ concentrations.

The measured $^{14}C$ concentrations reflect the dynamics of cell cycle activity, which is described by rates of cell birth and cell death [2, 22, 23, 24]. We defined a population balance equation for $^{14}C$ concentration structured cell populations, which we used to predict $^{14}C$ concentration dynamics from cell renewal rates. The cell renewal rates are allowed to change over time, which enables us to model different rates before disease onset, during the disease and after treatment with LVAD. We estimated the cell renewal rates using a Bayesian inference framework [22]. In this model, we assumed that new cardiomyocytes are generated by self-duplication of pre-existing cardiomyocytes [4, 7]. We accounted for the polyploidy of cardiomyocytes and accordingly defined eight populations by their nuclear ploidy level and their number of nuclei. We restricted the model parameters such that the model would correctly predict our here established ploidy and multinucleation time courses. Physiological polyploidy increases over the first two decades of life and plateaus in healthy subjects during adulthood [2] (Supplementary Methods, Figure B). As it remains unclear if all these cardiomyocyte populations can self-duplicate and how these populations can contribute to each other, we tested two principal scenarios: Scenario I allowed self-duplication of all populations, as well as increase to the next higher nuclear ploidy level and binucleation (Supplementary **Fig. S3A**). The more restricted scenario II only allowed self-duplication of mononucleated diploid cardiomyocytes and restricted the exchange between mono- and binucleated cardiomyocytes (Supplementary **Fig. S3B**). To validate these scenarios, we fitted $^{14}C$ data from healthy hearts reported by Bergmann et al. 2015. For both scenarios, we obtained similar goodness of fit and renewal rates of approximately 0.5% per year (Supplementary **Fig. S3C**). This is consistent with our previously published rates supporting the robustness of our approach with respect to specific model assumptions [2, 3]. As the restricted scenario II is in line with previous findings that diploid mononucleated cardiomyocytes contribute to cardiomyocyte renewal [25], we utilized this less complex scenario II for further analyses.

### Population dynamic

To reflect the different ploidy populations present in the adult human heart, we included mono- and binucleated cells with ploidy ranging from 2n up to 16n, resulting in 8 populations indexed by their nucleation and ploidy levels, i.e. $1x2n$, $1x4n$, $1x8n$, $1x16n$, $2x2n$, $2x4n$, $2x8n$, $2x16n$ (**Supplementary Fig. S3**). Here, in these supplementary methods we will use the term 2n cells to describe the combined 1x2n and 2x2n cell populations (and higher ploidy levels accordingly). We used our measurements of ploidy and binucleation, to constrain the number of cells ($N_{1x2n}$, $N_{1x4n}$,…) in each of the cell populations. We observed that ploidy levels are equally distributed in mono- and binucleated cardiomyocytes (**Fig. 1J**), i.e.

$$\frac{N_{1x2n}}{N_{2x2n}} \approx \frac{N_{1x4n}}{N_{2x4n}} \approx \frac{N_{1x8n}}{N_{2x8n}} \approx \frac{N_{1x16n}}{N_{2x16n}} = \frac{1-m(t)}{m(t)},$$

where $1-m(t)$ and $m(t)$ are the fractions of mononucleated and binucleated cardiomyocytes, respectively. Thus, we can write the number of cells in each cell population as:

$$\begin{aligned} N_{1xi}(t) &= f_i(t)(1-m(t))N_{\text{total}}(t), \quad && i \in \{2n, 4n, 8n, 16n\} \\ N_{2xi}(t) &= f_i(t)m(t)N_{\text{total}}(t), \quad && i \in \{2n, 4n, 8n, 16n\} \quad (1) \\ f_{2n}(t) + f_{4n}(t) &+ f_{8n}(t) + f_{16n}(t) = 1 && \end{aligned}$$

where $f_{2n}(t) = \frac{N_{1x2n}(t)+N_{2x2n}(t)}{N_{total}(t)}$ is the ploidy fraction for 2n cells (and higher ploidy levels accordingly).

### Model for the fractions of binucleated cells

For a healthy person, the mean fraction of binucleated cardiomyocytes is approximately 20% (**Fig. 1G**). We found that the fraction of binucleated cells does not depend on age (Bergmann et al., 2015) (**Supplementary Fig. A**). Consequently, we model the fraction of binucleated cells for healthy subjects to be constant, $m(t) = 20\%$. For subjects with diseases, the fraction of binucleated cells also does not depend on age and is approximately 35 % (**Supplementary Fig. S1D**). In our model, we assume a linear transition from the healthy state to the disease state starting at the time of its diagnosis:

$$m(t) = \begin{cases} 20\% & if\ t \leq t_d \\ 20\% - \frac{t + t_d}{t_c - t_d}\,(20\% - 35\%) & if\ t > t_d \end{cases}, \qquad (2)$$

where $t_c$ is the age of the subject at the collection date and $t_d$ the age of the subject at the onset of the disease i.e. the time of the diagnosis.

### Model for ploidy fractions

Ploidy levels are age-dependent: Adults show lower fractions of 2n cells and higher fractions of 4n and 8n cells than children (**Supplementary Fig. B**). For healthy subjects, we model the ploidy dynamics with an exponential function:

$$f_i^H(t) = (f_i^0 - g_i)\exp\left(-\frac{t}{\tau}\right) + \frac{f_i^d - f_i^0 \exp\left(-\frac{t_d}{\tau}\right)}{1 - \exp\left(-\frac{t_d}{\tau}\right)} \qquad (3)$$

The decay constant $\tau = 12$ years was estimated by performing a least square fit of the function $f_{2n}^H(t)$ to the ploidy fraction measurements of healthy subjects. Based on the limited data for children we use the following fraction at birth: $f_{2n}^0 = 0.955$, $f_{4n}^0 = 0.04$, $f_{8n}^0 = 0.005$ and $f_{16n}^0 =$

0. The fraction at the collection time $f_i^d$ is known from the measured values for each subject (**Supplementary Fig. A**).

For subjects with heart disease, we model the ploidy fraction dynamics with an exponential function until disease onset (similar to healthy as above) and with a linear function after disease onset (**Supplementary Fig. B**):

$$f_i(t) = \begin{cases} (f_i^0 - g_i)\exp\left(-\frac{t}{\tau}\right) + \frac{f_i^h - f_i^0 \exp\left(-\frac{t_d}{\tau}\right)}{1 - \exp\left(-\frac{t_d}{\tau}\right)} & if\ t \leq t_d \\ f_i^h - \frac{t - t_d}{t_c - t_d}\,(f_i^h - f_i^d) & if\ t > t_d \end{cases} \quad (4)$$

The fractions $f_i^0$, $f_i^h$ and $f_i^d$ correspond to the fraction at birth (superscript 0), at the onset of the disease (h) and at the collection date (d), respectively. The determination of the healthy fraction $f_i^h$ for subjects with heart disease is described in detail in the next section.

### Estimation of the ploidy fraction $f_i^h$ before onset of disease for subjects with heart disease

For subjects with heart disease, the ploidy fraction $f_i^h$ at the onset of the disease when the subject was healthy cannot be measured and hence needs to be estimated. To do so, we assumed that the ploidy fraction in subjects with heart disease would follow similar dynamics as the ploidy fraction in healthy subjects. For a given subject with heart disease, we assume disease onset at age of diagnosis, $t_d$. For each healthy subject, we can predict the ploidy fraction at time $t_d$ using $f_i^H(t_d)$ from equation (3). Because all subjects in this study were diagnosed during adulthood, we can approximate $f_i^H(t_d) \approx f_i^d$, which is the ploidy fraction that was measured for each subject. In a first approach, we estimated the ploidy fraction $f_i^h$ at the onset by taking the mean of $f_i^d$ over all healthy subjects. However, this led to a situation where the model was impossible to solve (or only with negative turnover rates, which are of course not possible).

Hence, we refined our approach to estimate $f_i^h$ in the following way, by taking the variability in the healthy population into account: We use the average fractions over all subjects for $f_{8n}^d$ and $f_{16n}^d$ for the 8n ($f_{8n}^h = 7\%$) and 16n ($f_{16n}^h$=0%) cell populations, respectively. Since the sum of all fractions is 1, only the fraction $f_{2n}^h$ needs to be estimated and $f_{4n}^h$ can be computed from equations (1). We estimate $f_{2n}^h$ using a Baysian approach: As the prior probability distribution we use a Gaussian that is parameterized with the mean and variance of $f_i^d$ over all subjects. We set the likelihood of $f_{2n}^h$ to 1 if the model can be solved for that value and 0 otherwise. The posterior probability distribution of $f_{2n}^h$ is then given by a truncated Normal distribution. We then chose the median of this posterior distribution as an estimate for the $f_i^h$.

### $^{14}$C concentration model for scenario I

We model the state of each cell population $i$ at time $t$ with $n_i(t, c)$, which is the density of cells with genomic $^{14}$C concentration $c$. In each cell population cell death happens at rate $\delta_i(t)$, mitotic cell division happens at rate $\beta_i(t)$, and non-productive cell cycle activities that transport cells from population $i$ to $j$ (e.g. from 2n to 4n) happen at rates $\kappa_{i,j}(t)$. In the following, we will omit the time-dependence of the rates in our notation, i.e. we write $\beta_i$ instead of $\beta_i(t)$. Then, we model the cell density dynamics in each cell population $i$ with an equation of the form

$$\frac{\partial n_i(t,c)}{\partial t} = 2\,\beta_i \int dc' f_{\beta_i}\big(c,c' \big| c_a(t+b)\big) n_i(t,c') + \sum_j \kappa_{j,i} \int dc' f_{\kappa_{i,j}}\big(c,c' \big| c_a(t+b)\big) n_j(t,c')$$

$$- \left( \beta_i + \delta_i + \sum_j \kappa_{i,j} \right) n(t,c)$$

where the first term models the gain due to cell divisions, the second term models gain due to non-productive cell cycle activity in other cell populations and the third term models the loss due to divisions, death and non-productive cell cycle activity. $b$ is the birth date of the subject and $c_a(t+b)$ the atmospheric $^{14}C$ concentration shifted by one year due to the food chain delay. The kernel $f_{\beta_i}\big(c,c' \big| c_a(t+b)\big)$ describes the probability of a cell with $^{14}C$ concentration $c'$ to divide into a daughter cell with $^{14}C$ concentration c which depends on the food chain delayed atmospheric $^{14}C$ concentration at the time of the division. The average $^{14}C$ concentration of each pair of daughter cells must be $c_m = \frac{c' + c_{a(t+b)}}{2}$. So, the kernels must be symmetric with respect to $c_m$:

$$f_{\beta_i}\big(x + c_m, c' \big| c_a(t+b)\big) = f_{\beta_i}\big(-x + c_m, c' \big| c_a(t+b)\big) \quad \forall x$$

Furthermore, we require the kernel to be cell number conserving:

$$\int_0^\infty dc \;\; f_{\beta_i}\big(c,c' \big| c_a(t+b)\big) = 1$$

And the average $^{14}C$ concentration must be:

$$\int_0^\infty dc \;\; c\, f_{\beta_i}\big(c,c' \big| c_a(t+b)\big) = c_m = \frac{c' + c_{a(t+b)}}{2}.$$

For cardiomyocytes, we are using 8 populations and assume that each population has the same death rate ($\delta_i = \delta$), but different successful ($\beta_i$) and unsuccessful ($\kappa_{i,j}$) division rates. Unsuccessful divisions can either increase ploidy level by one or they can turn mononucleated cells into binucleated cells. For every unsuccessful division, the resulting cell contains the DNA from the mother cell as well as the newly synthesized DNA, resulting in an average $^{14}$C concentration between the food chain delayed atmospheric $^{14}$C concentration and the one of the mother cell. Hence, the kernel is a delta distribution:

$$f_{\kappa_{i,j}}\left(c, c' \middle| c_a(t+b)\right) = \delta\left(c - \frac{c' + c_{a(t+b)}}{2}\right).$$

The integral over $c'$ can be solved directly:

$$\int dc' f_\kappa\left(c, c' \middle| c_a(t+b)\right) n_i(t, c') = n_i\big(t, 2\,c - c_a(t+b)\big).$$

Now, we can write down the equations for the full mode. For more readable equations, the arguments for the kernels and atmospheric $^{14}$C concentration are omitted.

$$
\begin{aligned}
\frac{\partial c_{1x2n}(c,t)}{\partial t} &= -(\beta_{1x2n} + \delta + \kappa_{1x2n,1x4n} + \kappa_{1x2n,2x2n})c_{1x2n}(c,t) + 2\beta_{1x2n}\int dc' f_\beta c_{1x2n}(c',t), \\
\frac{\partial c_{1x4n}(c,t)}{\partial t} &= -(\beta_{1x4n} + \delta + \kappa_{1x4n,1x8n} + \kappa_{1x4n,2x4n})c_{1x4n}(c,t) \\
&\quad +2\beta_{1x4n}\int dc' f_\beta c_{1x4n}(c',t) + \kappa_{1x2n,1x4n}c_{1x2n}(2c - c_a, t), \\
\frac{\partial c_{1x8n}(c,t)}{\partial t} &= -(\beta_{1x8n} + \delta + \kappa_{1x8n,1x16n} + \kappa_{1x8n,2x8n})c_{1x8n}(c,t) \\
&\quad +2\beta_{1x8n}\int dc' f_\beta c_{1x8n}(c',t) + \kappa_{1x4n,1x8n}c_{1x4n}(2c - c_a, t), \\
\frac{\partial c_{1x16n}(c,t)}{\partial t} &= -(\beta_{1x16n} + \delta + \kappa_{1x16n,2x16n})c_{1x16n}(c,t) \\
&\quad +2\beta_{1x16n}\int dc' f_\beta c_{1x16n}(c',t) + \kappa_{1x8n,1x16n}c_{1x8n}(2c - c_a, t), \\
\frac{\partial c_{2x2n}(c,t)}{\partial t} &= -(\beta_{2x2n} + \delta + \kappa_{2x2n,2x4n})c_{2x2n}(c,t) \\
&\quad +2\beta_{2x2n}\int dc' f_\beta c_{2x2n}(c',t) + \kappa_{1x2n,2x2n}c_{1x2n}(2c - c_a, t), \\
\frac{\partial c_{2x4n}(c,t)}{\partial t} &= -(\beta_{2x4n} + \delta + \kappa_{2x4n,2x8n})c_{2x4n}(c,t) \\
&\quad +2\beta_{2x4n}\int dc' f_\beta c_{2x4n}(c',t) + \kappa_{2x2n,2x4n}c_{2x2n}(2c - c_a, t) + \kappa_{1x4n,2x4n}c_{1x4n}(2c - c_a, t), \\
\frac{\partial c_{2x8n}(c,t)}{\partial t} &= -(\beta_{2x8n} + \delta + \kappa_{2x8n,2x16n})c_{2x8n}(c,t) \\
&\quad +2\beta_{2x8n}\int dc' f_\beta c_{2x8n}(c',t) + \kappa_{2x4n,2x8n}c_{2x4n}(2c - c_a, t) + \kappa_{1x8n,2x8n}c_{1x8n}(2c - c_a, t), \\
\frac{\partial c_{2x16n}(c,t)}{\partial t} &= -(\beta_{2x16n} + \delta)c_{2x16n}(c,t) + 2\beta_{2x16n}\int dc' f_\beta c_{2x16n}(c',t) \\
&\quad +\kappa_{2x8n,2x16n}c_{2x8n}(2c - c_a, t) + \kappa_{1x16n,2x16n}c_{1x16n}(2c - c_a, t)
\end{aligned}
\tag{5}
$$

Integrating equations (5) over all $^{14}C$ concentration results in total number of cells per population $N_i(t) = \int_0^\infty dc n_i(t,c)$ and their dynamic are:

$$
\begin{aligned}
\frac{\partial N_{1x2n}(t)}{\partial t} &= 2\beta_{1x2n}N_{1x2n}(t) + (-\beta_{1x2n} - \delta - \kappa_{1x2n,2x2n} - \kappa_{1x2n,1x4n})N_{1x2n}(t) \\
\frac{\partial N_{1x4n}(t)}{\partial t} &= \kappa_{1x2n,1x4n}N_{1x2n}(t) + 2\beta_{1x4n}N_{1x4n}(t) + (-\beta_{1x4n} - \delta - \kappa_{1x4n,2x4n} - \kappa_{1x4n,1x8n})N_{1x4n}(t) \\
\frac{\partial N_{1x8n}(t)}{\partial t} &= \kappa_{1x4n,1x8n}N_{1x4n}(t) + 2\beta_{1x8n}N_{1x8n}(t) + (-\beta_{1x8n} - \delta - \kappa_{1x8n,1x16n} - \kappa_{1x8n,2x8n})N_{1x8n}(t) \\
\frac{\partial N_{1x16n}(t)}{\partial t} &= 2\beta_{1x16n}N_{1x16n}(t) + (-\beta_{1x16n} - \delta - \kappa_{1x16n,2x16n})N_{1x16n}(t) + \kappa_{1x8n,1x16n}N_{1x8n}(t) \\
\frac{\partial N_{2x2n}(t)}{\partial t} &= \kappa_{1x2n,2x2n}N_{1x2n}(t) + 2\beta_{2x2n}N_{2x2n}(t) + (-\beta_{2x2n} - \delta - \kappa_{2x2n,2x4n})N_{2x2n}(t) \\
\frac{\partial N_{2x4n}(t)}{\partial t} &= \kappa_{2x2n,2x4n}N_{2x2n}(t) + 2\beta_{2x4n}N_{2x4n}(t) + \left(-\beta_{2x4n} - \delta - \kappa_{2x4n,2x8n}\right)N_{2x4n}(t) \\
&\quad + \kappa_{1x4n,2x4n}N_{1x4n}(t) \\
\frac{\partial N_{2x8n}(t)}{\partial t} &= \kappa_{2x4n,2x8n}N_{2x4n}(t) + 2\beta_{2x8n}N_{2x8n}(t) + \left(-\beta_{2x8n} - \delta - \kappa_{2x8n,2x16n}\right)N_{2x8n}(t) \\
&\quad + \kappa_{1x8n,2x8n}N_{1x8n}(t) \\
\frac{\partial N_{2x16n}(t)}{\partial t} &= \kappa_{1x16n,2x16n}N_{1x16n}(t) + 2\beta_{2x16n}N_{2x16n}(t) + (-\beta_{2x16n} - \delta)N_{2x16n}(t) \\
&\quad + \kappa_{2x8n,2x16n}N_{2x8n}(t)
\end{aligned}
\tag{6}
$$

Finally, the mean $^{14}C$ concentration are given by $\bar{c}_i(t) = \frac{\int_0^\infty dc\, c\, n_i(t,c)}{N_i(t)}$ and applying this to equations (5) leads to:

$$
\begin{aligned}
\frac{\partial \overline{c_{1x2n}}(t)}{\partial t} &= \beta_{1x2n}(-\overline{c_{1x2n}}(t) + c_a) \\
\frac{\partial \overline{c_{1x4n}}(t)}{\partial t} &= \beta_{1x4n}(-\overline{c_{1x4n}}(t) + c_a) + \frac{\kappa_{1x2n,1x4n}(\overline{c_{1x2n}}(t) - 2\overline{c_{1x4n}}(t) + c_a)N_{1x2n}(t)}{2N_{1x4n}(t)} \\
\frac{\partial \overline{c_{1x8n}}(t)}{\partial t} &= \beta_{1x8n}(-\overline{c_{1x8n}}(t) + c_a) + \frac{\kappa_{1x4n,1x8n}(\overline{c_{1x4n}}(t) - 2\overline{c_{1x8n}}(t) + c_a)N_{1x4n}(t)}{2N_{1x8n}(t)} \\
\frac{\partial \overline{c_{1x16n}}(t)}{\partial t} &= \frac{2\beta_{1x16n}(-\overline{c_{1x16n}}(t) + c_a)N_{1x16n}(t) + \kappa_{1x8n,1x16n}(-2\overline{c_{1x16n}}(t) + \overline{c_{1x8n}}(t) + c_a)N_{1x8n}(t)}{2N_{1x16n}(t)} \\
\frac{\partial \overline{c_{2x2n}}(t)}{\partial t} &= \beta_{2x2n}(-\overline{c_{2x2n}}(t) + c_a) + \frac{\kappa_{1x2n,2x2n}(\overline{c_{1x2n}}(t) - 2\overline{c_{2x2n}}(t) + c_a)N_{1x2n}(t)}{2N_{2x2n}(t)} \\
\frac{\partial \overline{c_{2x4n}}(t)}{\partial t} &= \frac{\kappa_{2x2n,2x4n}\overline{c_{2x2n}}(t)N_{2x2n}(t) + c_a\left(\kappa_{2x2n,2x4n}N_{2x2n}(t) + 2\beta_{2x4n}N_{2x4n}(t)\right)}{2N_{2x4n}(t)} \\
&+ \frac{\kappa_{1x4n,2x4n}(\overline{c_{1x4n}}(t) + c_a)N_{1x4n}(t)}{2N_{2x4n}(t)} \\
&- \frac{2\overline{c_{2x4n}}(t)\left(\kappa_{2x2n,2x4n}N_{2x2n}(t) + \beta_{2x4n}N_{2x4n}(t) + \kappa_{1x4n,2x4n}N_{1x4n}(t)\right)}{2N_{2x4n}(t)} \qquad (7) \\
\frac{\partial \overline{c_{2x8n}}(t)}{\partial t} &= \frac{\kappa_{2x4n,2x8n}\overline{c_{2x4n}}(t)N_{2x4n}(t) + c_a\left(\kappa_{2x4n,2x8n}N_{2x4n}(t) + 2\beta_{2x8n}N_{2x8n}(t)\right)}{2N_{2x8n}(t)} \\
&+ \frac{\kappa_{1x8n,2x8n}(\overline{c_{1x8n}}(t) + c_a)N_{1x8n}(t)}{2N_{2x8n}(t)} \\
&- \frac{2\overline{c_{2x8n}}(t)\left(\kappa_{2x4n,2x8n}N_{2x4n}(t) + \beta_{2x8n}N_{2x8n}(t) + \kappa_{1x8n,2x8n}N_{1x8n}(t)\right)}{2N_{2x8n}(t)} \\
\frac{\partial \overline{c_{2x16n}}(t)}{\partial t} &= \frac{\kappa_{1x16n,2x16n}\overline{c_{1x16n}}(t)N_{1x16n}(t) + c_a\left(\kappa_{1x16n,2x16n}N_{1x16n}(t) + 2\beta_{2x16n}N_{2x16n}(t)\right)}{2N_{2x16n}(t)} \\
&+ \frac{\kappa_{2x8n,2x16n}(\overline{c_{2x8n}}(t) + c_a)N_{2x8n}(t)}{2N_{2x16n}(t)} \\
&- \frac{2\overline{c_{2x16n}}(t)\left(\kappa_{1x16n,2x16n}N_{1x16n}(t) + \beta_{2x16n}N_{2x16n}(t) + \kappa_{2x8n,2x16n}N_{2x8n}(t)\right)}{2N_{2x16n}(t)}
\end{aligned}
$$

Next, we use the 8 constraints on ploidy-level and binucleation given by equations (1) to reduce the number of free parameters by 8. Solving equations (7) for the following parameters leads to:

$$
\begin{aligned}
\beta_{1x2n} &= \frac{(-\beta_{1x16n}+\delta)N_{1x16n}(t)-\beta_{2x16n}N_{2x16n}(t)-\beta_{2x2n}N_{2x2n}(t)-\beta_{2x4n}N_{2x4n}(t)}{N_{1x2n}(t)} \\
&-\frac{\beta_{2x8n}N_{2x8n}(t)}{N_{1x2n}(t)}-\frac{\beta_{1x4n}N_{1x4n}(t)}{N_{1x2n}(t)}-\frac{\beta_{1x8n}N_{1x8n}(t)}{N_{1x2n}(t)} \\
&+\frac{\delta\big(N_{1x2n}(t)+N_{2x16n}(t)+N_{2x2n}(t)+N_{2x4n}(t)+N_{2x8n}(t)+N_{1x4n}(t)+N_{1x8n}(t)\big)+\partial_t N_{1x16n}(t)}{N_{1x2n}(t)} \\
&+\frac{\partial_t N_{1x2n}(t)+\partial_t N_{2x16n}(t)+\partial_t N_{2x2n}(t)+\partial_t N_{2x4n}(t)+\partial_t N_{2x8n}(t)+\partial_t N_{1x4n}(t)+\partial_t N_{1x8n}(t)}{N_{1x2n}(t)} \\
\kappa_{1x2n,1x4n} &= \frac{\big(-\beta_{1x16n}+\delta+\kappa_{1x16n,2x16n}\big)N_{1x16n}(t)+\big(-\beta_{1x4n}+\delta+\kappa_{1x4n,2x4n}\big)N_{1x4n}(t)}{N_{1x2n}(t)} \\
&+\frac{\big(-\beta_{1x8n}+\delta+\kappa_{1x8n,2x8n}\big)N_{1x8n}(t)+\partial_t N_{1x16n}(t)+\partial_t N_{1x4n}(t)+\partial_t N_{1x8n}(t)}{N_{1x2n}(t)} \\
\kappa_{1x4n,1x8n} &= \frac{\big(-\beta_{1x16n}+\delta+\kappa_{1x16n,2x16n}\big)N_{1x16n}(t)+\big(-\beta_{1x8n}+\delta+\kappa_{1x8n,2x8n}\big)N_{1x8n}(t)}{N_{1x4n}(t)} \\
&+\frac{\partial_t N_{1x16n}(t)+\partial_t N_{1x8n}(t)}{N_{1x4n}(t)} \\
\kappa_{1x8n,1x16n} &= \frac{\big(-\beta_{1x16n}+\delta+\kappa_{1x16n,2x16n}\big)N_{1x16n}(t)+\partial_t N_{1x16n}(t)}{N_{1x8n}(t)} \\
\kappa_{1x2n,2x2n} &= \frac{-\kappa_{1x16n,2x16n}N_{1x16n}(t)+(-\beta_{2x16n}+\delta)N_{2x16n}(t)-\beta_{2x2n}N_{2x2n}(t)-\beta_{2x4n}N_{2x4n}(t)}{N_{1x2n}(t)} \\
&-\frac{\beta_{2x8n}N_{2x8n}(t)}{N_{1x2n}(t)}+\frac{\delta\big(N_{2x2n}(t)+N_{2x4n}(t)+N_{2x8n}(t)\big)-\kappa_{1x4n,2x4n}N_{1x4n}(t)}{N_{1x2n}(t)} \\
&-\frac{\kappa_{1x8n,2x8n}N_{1x8n}(t)}{N_{1x2n}(t)}+\frac{\partial_t N_{2x16n}(t)+\partial_t N_{2x2n}(t)+\partial_t N_{2x4n}(t)+\partial_t N_{2x8n}(t)}{N_{1x2n}(t)} \\
\kappa_{2x2n,2x4n} &= \frac{-\kappa_{1x16n,2x16n}N_{1x16n}(t)+(-\beta_{2x16n}+\delta)N_{2x16n}(t)+(-\beta_{2x4n}+\delta)N_{2x4n}(t)}{N_{2x2n}(t)} \\
&+\frac{(-\beta_{2x8n}+\delta)N_{2x8n}(t)-\kappa_{1x4n,2x4n}N_{1x4n}(t)-\kappa_{1x8n,2x8n}N_{1x8n}(t)+\partial_t N_{2x16n}(t)}{N_{2x2n}(t)} \\
&+\frac{\partial_t N_{2x4n}(t)+\partial_t N_{2x8n}(t)}{N_{2x2n}(t)} \\
\kappa_{2x4n,2x8n} &= \frac{-\kappa_{1x16n,2x16n}N_{1x16n}(t)+(-\beta_{2x16n}+\delta)N_{2x16n}(t)+(-\beta_{2x8n}+\delta)N_{2x8n}(t)-\kappa_{1x8n,2x8n}N_{1x8n}(t)}{N_{2x4n}(t)} \\
&+\frac{\partial_t N_{2x16n}(t)+\partial_t N_{2x8n}(t)}{N_{2x4n}(t)} \\
\kappa_{2x8n,2x16n} &= \frac{-\kappa_{1x16n,2x16n}N_{1x16n}(t)+(-\beta_{2x16n}+\delta)N_{2x16n}(t)+\partial_t N_{2x16n}(t)}{N_{2x8n}(t)}
\end{aligned}
\tag{8}
$$

### Scenario II model

For this model, we assume that only the mononucleated 2n population can self-renew i.e. all $\beta_i$ are set to zero except for $\beta_{1x2n}$. Furthermore, we assume polyploid cells cannot increase their nucleation resulting in $\kappa_{1x4n,2x4n} = \kappa_{1x8n,2x8n} = \kappa_{1x16n,2x16n} = 0$. Thus, the scenario II model has only the death rate $\delta$ as a free parameter.

### 2-Phase model for diseased subjects

For the diseased subjects, we extend the scenario II model with a time-dependent death rate: $\delta(t) = \begin{cases} \delta_h \; if \; t \leq a_d \\ \delta_d \; if \; t > a_d \end{cases}$, where $\delta_h$ is the death rate when the subject was healthy, $\delta_h$the one when the subject was ill and $a_d$ is the age of the subjects when the disease was diagnosed. For the healthy phase, we assume the same behavior as for the healthy subjects. Thus, the result from the healthy subjects defines the death rate $\delta_h$, leaving only the death rate $\delta_d$ as a free parameter.

### Parameter estimation with Bayesian inference

We measure the mean $^{14}$C concentration of cardiomyocyte nuclei without differentiating the ploidy or nucleation. The predicted mean $^{14}$C concentration from our model is:

$$\bar{c}(t) = \frac{\sum \mathrm{w_i}\, \mathrm{N_i(t)}\; \bar{c}_\mathrm{i}(t)}{\sum \mathrm{w_i}\, \mathrm{N_i(t)}},$$

where $\mathrm{w_i}$ is the number of nuclei of population i.

We used an additive Gaussian noise model to define a likelihood for our data with a sample independent variance of σ. We used uniform priors in the log space for the unknown rate parameters

$$\mathcal{U}(\log_{10} 10^{-6}\mathrm{years}, \log_{10} 10^{0}\mathrm{years})$$

and a uniform prior for σ:

$$\sigma \sim \mathcal{U}(0, 0.5).$$

Due to the cell number constraints, we must eliminate one parameter for each population. Mathematically, these rates can become negative. However, because a negative rate is physically not possible, an infinitely unlikely value must be defined for the likelihood. For such cases, the likelihood is set to $-\infty$.

To numerically solve the Bayesian inference problem (Fröhlich et al., 2017; Hasenauer et al., 2012; Virtanen et al., 2020), we used Markov chain Monte Carlo (MCMC) sampling as implemented in emcee: The MCMC Hammer (Foreman-Mackey et al., 2013). For each scenario, we used 3000 samples, of which 1000 were used for the burn-in phase, and the number of chains was 50 times the number of parameters resulting in 3000*50*number of parameters samples. The initial values for the unknown parameters were drawn from the priors. However, if a set of initial values results in a negative rate it is discarded, and a new sample is drawn. This avoids "stuck" chains in an area with a flat likelihood of $-\infty$. To evaluate the predictive power (goodness of the fit) of the model and compare them, we utilize the leave-one-out cross validation (LOO). The LOO is estimated with the Pareto-smoothed importance sampling (Vehtari et al., 2017).

### Results

The numerical solution of the Bayesian inference yields the approximated posterior distributions for the estimated parameters. These posterior distributions are depicted in **Supplementary Fig. C, D** for the healthy cases and in **Supplementary Fig. E-I** for the diseased cases. The point estimates and the model comparison for the healthy subjects are summarized in **Table A**. The point estimates for the diseased subjects fitted with the 2-phase model are summarized in **Table B**.

### Model selection for healthy subjects

The model comparison shows that scenarios I and II have similar predictive power (see **Table A**). Given that the populations dynamics are the same in all models and the death rate is the only process for cell death, it corresponds to the turnover rate. In the margins of error, the death rate from the scenarios I and II are identical. Thus, the additional processes of the scenario I model have no impact on the turnover rate of the tissue. This indicates that scenario II is sufficient to explain the data and the additional processes of the scenario I can be neglected.

Individually, the additional processes from scenario I can have rates larger than zero. To evaluate whether these processes are important we calculate their contribution to the overall synthesized DNA. Since the amount of synthesized DNA depends on the age of the subjects, we estimate an upper boundary by choosing the age for each subject where the contribution from these additional processes to the synthesized DNA is the largest. Then we use the average over all subjects. To include the information of the posterior distributions, we perform this calculation for 100,000 sets of parameters from the MCMC sampling and then take the median for the point estimates. The additional processes of the scenario I contribute at most 11% to the total DNA synthesis.

| MODEL | LOO | WEIGHT | PARAMETER | VALUE | CONFIDENCE INTERVAL |
|---|---|---|---|---|---|
| **SCENARIO II** | **32.55** | **50%** | $\delta$ | 0.55%/year | [0.44%/year - 0.71%/year] |
| | | | $\sigma$ | 0.038 | [0.030 - 0.043] |
| **SCENARIO I** | **32.53** | **50%** | $\delta$ | 0.57%/year | [0.46%/year - 0.74%/year] |
| | | | $\kappa_{1x4n,2x4n}$ | 0.003%/year | [0%/year - 0.03%/year] |
| | | | $\kappa_{1x8n,2x8n}$ | 0.00347%/year | [0%/year - 0.03%/year] |
| | | | $\kappa_{1x16n,2x16n}$ | 0.1%/year | [0.01%/year - 100%/year] |
| | | | $\beta_{1x4n}$ | 0.008%/year | [0%/year - 0.2%/year] |

| | | | |
|---|---|---|---|
| | $\beta_{1x8n}$ | 0.008%/year | [0%/year - 0.3%/year] |
| | $\beta_{1x16n}$ | 0.07%/year | [0%/year - 0.8%/year] |
| | $\beta_{2x2n}$ | 0.01%/year | [0.001%/year - 0.3%/year] |
| | $\beta_{2x4n}$ | 0.007%/year | [0%/year - 0.04%/year] |
| | $\beta_{2x8n}$ | 0.007%/year | [0%/year - 0.1%/year] |
| | $\beta_{2x16n}$ | 0.08%/year | [0%/year - 1%/year] |
| | σ | 0.038 | [0.03 - 0.044] |

**Table A: Model selection and parameter estimates for the different models which are fitted using the healthy subjects.** LOO: value of the leave-one-out information criterion; lower values indicate higher predictive power (best model). Weights indicate the probability of each scenario being the best model among the tested scenarios.

### Results for diseased subjects

We use scenario II for the analysis of the diseased subjects because the added complexity of the scenario I leads only to unidentifiable parameters and has a neglectable impact on the turnover rate.

| GROUPS | PARAMETER | VALUE | CONFIDENCE INTERVAL |
|---|---|---|---|
| **NOLVAD/ICM+NICM** | $\delta_d$ | 0.03%/year | [0.002%/year - 2%/year] |
| | σ | 0.059 | [0.049 - 0.067] |
| noLVAD/**ICM** | $\delta_d$ | 0.01%/year | [0%/year - 0.1%/year] |
| | σ | 0.039 | [0.026 - 0.048] |
| noLVAD/**NICM** | $\delta_d$ | 0.03%/year | [0.002%/year - 2.6%/year] |
| | σ | 0.055 | [0.048 - 0.081] |

| | | | |
|---|---|---|---|
| **LVAD/RESPONDER** | $\delta_d$ | 3.0%/year | [2.1%/year – 5.0%/year] |
| | σ | 0.041 | [0.032 - 0.048] |
| **LVAD/NON-RESPONDER** | $\delta_d$ | 0.02%/year | [0%/year - 0.2%/year] |
| | σ | 0.070 | [0.053 - 0.083] |

**Table B: Parameter estimates for the 2-phase model which are fitted using the diseased subjects.**

**Figure A: Fraction of binucleated cardiomyocytes.** The linear regressions of the healthy subjects show no age-dependency (Slope=-0.03%/year ± 0.06%/year).

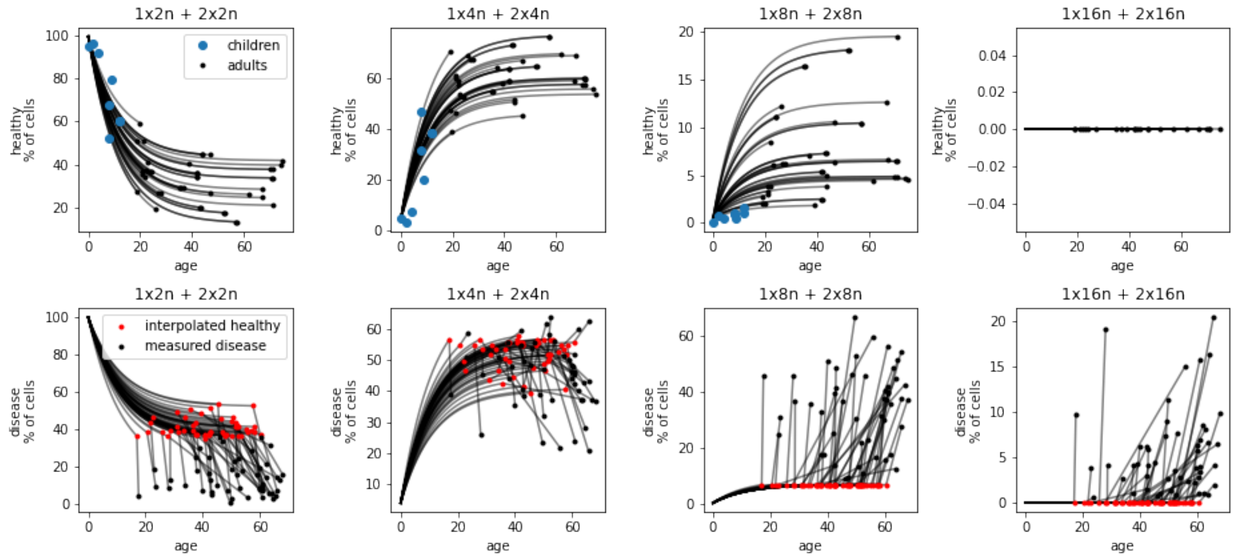


**Figure B: Age-dependent ploidy fractions.** The top row shows the ploidy levels for healthy subjects and the bottom row the ploidy fractions for the subjects with heart disease. The black dots depict the measured values for adults and the blue dots show the measured values for children (Bergmann et al 2015). The black lines visualize the results from our ploidy model (see equation (3)). The red dots show the interpolated ploidy fractions at the age of the onset of the disease.

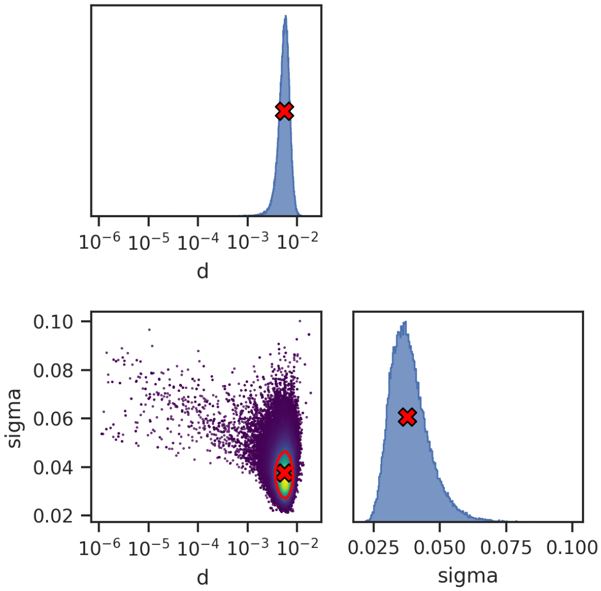


**Figure C: Posterior distributions of the scenario II with the healthy subjects estimated by MCMC sampling.** On diagonal panels: Marginal distributions of the model parameters. Off-diagonal panels: Marginal distributions of parameter pairs. Each dot is a sample from the MCMC chain, and the dots are color coded according to the local probability density. The 1-σ confidence region (CR) is enclosed by the red contour and contains the true parameter with 68% probability.

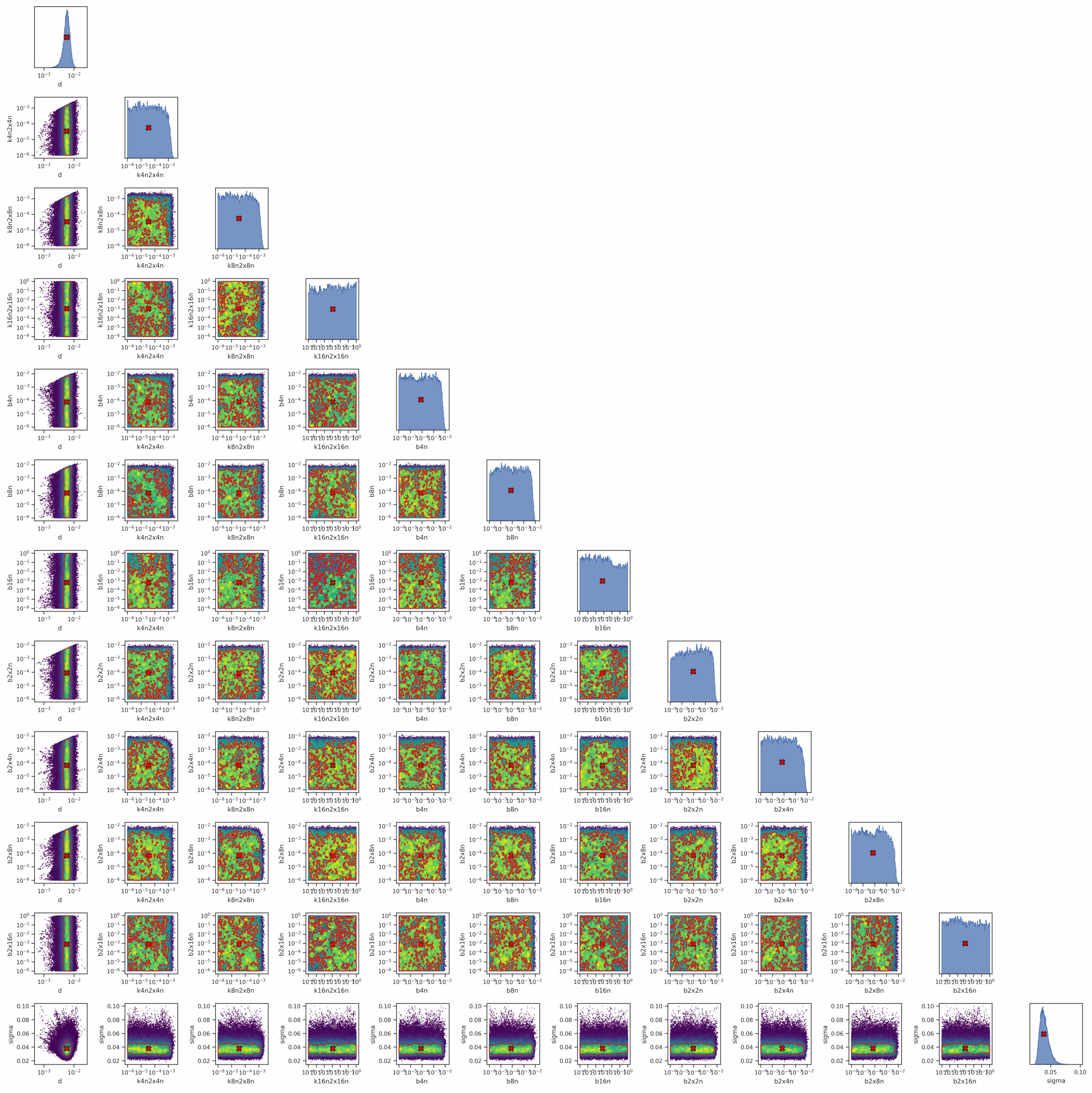


**Figure D: Posterior distributions of the scenario I with the healthy subjects estimated by MCMC sampling.** On diagonal panels: Marginal distributions of the model parameters. Off-diagonal panels: Marginal distributions of parameter pairs. Each dot is a sample from the MCMC chain, and the dots are color coded according to the local probability density. The 1-σ confidence region (CR) is enclosed by the red contour and contains the true parameter with 68% probability.

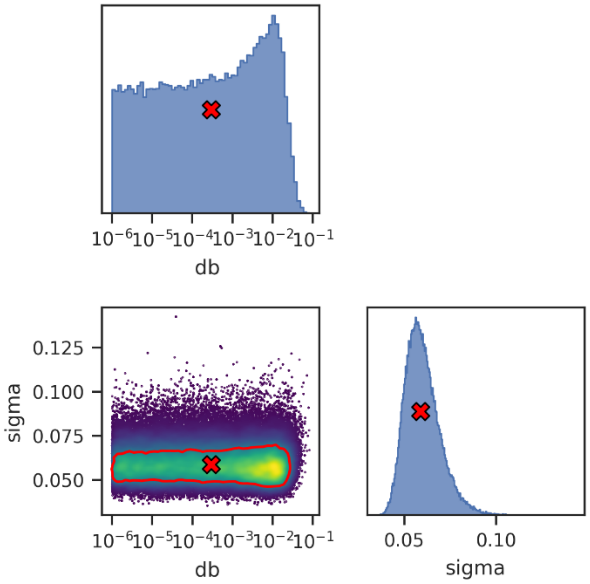


**Figure E: Posterior distributions of the 2-phase model with the diseased noLVAD/ ICM+NICM subjects estimated by MCMC sampling.** On diagonal panels: Marginal distributions of the model parameters. Off-diagonal panels: Marginal distributions of parameter pairs. Each dot is a sample from the MCMC chain, and the dots are color coded according to the local probability density. The 1-σ confidence region (CR) is enclosed by the red contour and contains the true parameter with 68% probability.

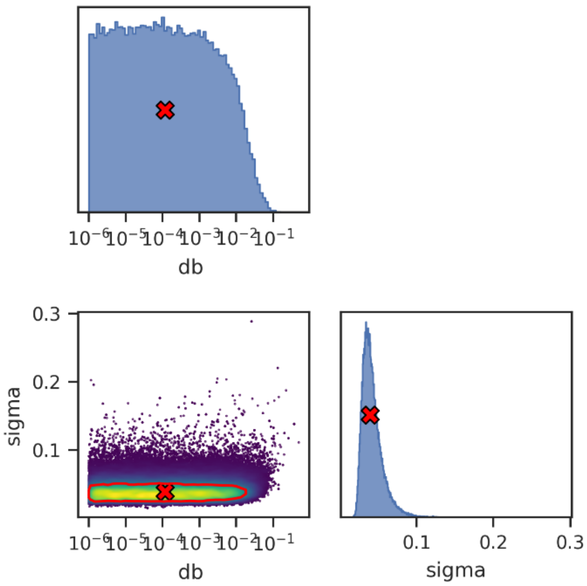


**Figure F: Posterior distributions of the 2-phase model with the noLVAD/ICM subjects estimated by MCMC sampling.** On diagonal panels: Marginal distributions of the model parameters. Off-diagonal panels: Marginal distributions of parameter pairs. Each dot is a sample from the MCMC chain, and the dots are color coded according to the local probability density. The 1-σ confidence region (CR) is enclosed by the red contour and contains the true parameter with 68% probability.

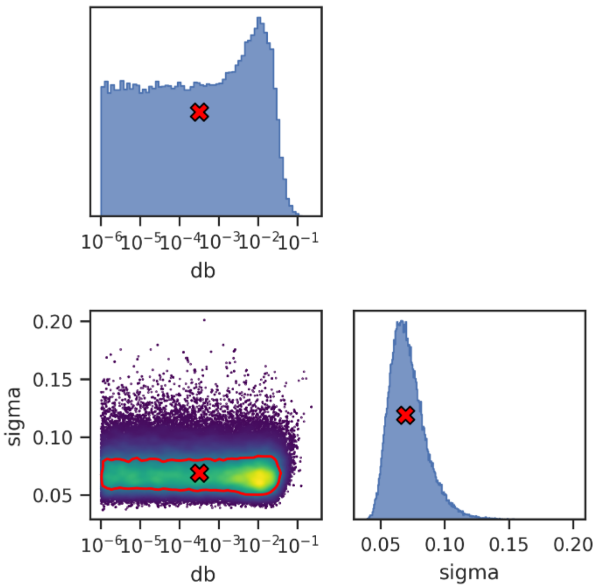


**Figure G: Posterior distributions of the 2-phase model with the noLVAD/NICM subjects estimated by MCMC sampling.** On diagonal panels: Marginal distributions of the model parameters. Off-diagonal panels: Marginal distributions of parameter pairs. Each dot is a sample from the MCMC chain, and the dots are color coded according to the local probability density. The 1-σ confidence region (CR) is enclosed by the red contour and contains the true parameter with 68% probability.

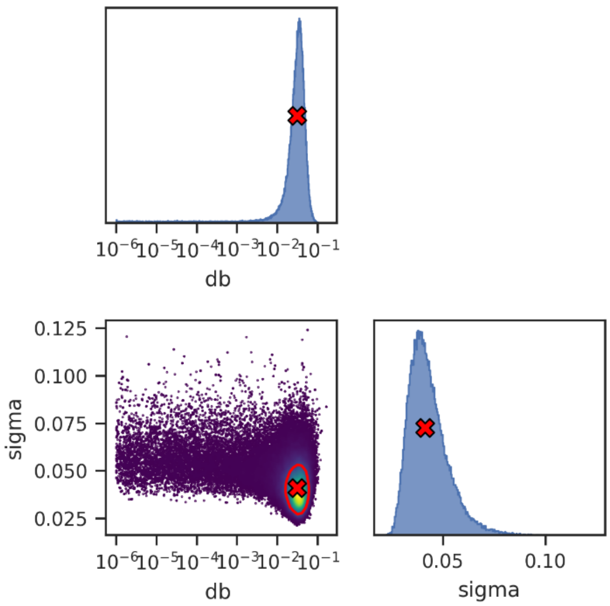


**Figure H: Posterior distributions of the 2-phase model with the LVAD/responder subjects estimated by MCMC sampling.** On diagonal panels: Marginal distributions of the model parameters. Off-diagonal panels: Marginal distributions of parameter pairs. Each dot is a sample from the MCMC chain, and the dots are color coded according to the local probability density. The 1-σ confidence region (CR) is enclosed by the red contour and contains the true parameter with 68% probability.

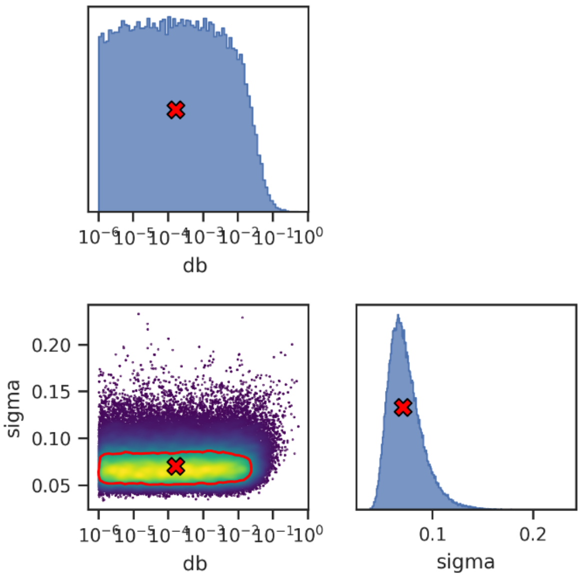


**Figure I: Posterior distributions of the 2-phase model with the LVAD/non-responder subjects estimated by MCMC sampling.** On diagonal panels: Marginal distributions of the model parameters. Off-diagonal panels: Marginal distributions of parameter pairs. Each dot is a sample from the MCMC chain, and the dots are color coded according to the local probability density. The 1-σ confidence region (CR) is enclosed by the red contour and contains the true parameter with 68% probability.
