## Supplemental Table 1 for "A latent cardiomyocyte regeneration potential in human heart disease"

Table 1. Patient characteristics of the study population for ^14^C analysis

| **Variable** | **Advanced HF n=24** | **LVAD non-Responder n=13** | **LVAD Responders n=15** |
| --- | --- | --- | --- |
| Male sex, n | 15 (63%) | 11 (85%) | 11 (73%) |
| Age at collection | 49.3 | 57.5 | 49.5 |
| Duration of heart failure symptoms (months) | 97.1 | 89.3 | 89.7 |
| Duration of LVAD (months) | N/A | 19.3 | 13.7 |
| Heart failure pathogenesis, n | | | |
| Ischemic cardiomyopathy | 8 (33%) | 6 (46%) | 2 (13%) |
| Nonischemic cardiomyopathy | 16 (66%) | 7 (54%) | 13 (87%) |
| Absolute LVEF change after LVAD, % | N/A | -4.2 | 13.7 |
